# *P. falciparum* K13 mutations present varying degrees of artemisinin resistance and reduced fitness in African parasites

**DOI:** 10.1101/2021.01.27.428390

**Authors:** Barbara H. Stokes, Kelly Rubiano, Satish K. Dhingra, Sachel Mok, Judith Straimer, Nina F. Gnädig, Jade R. Bath, Ioanna Deni, Kurt E. Ward, Josefine Striepen, Tomas Yeo, Leila S. Ross, Eric Legrand, Frédéric Ariey, Clark H. Cunningham, Issa M. Souleymane, Adama Gansané, Romaric Nzoumbou-Boko, Claudette Ndayikunda, Abdunoor M. Kabanywanyi, Aline Uwimana, Samuel J. Smith, Olimatou Kolley, Mathieu Ndounga, Marian Warsame, Rithea Leang, François Nosten, Timothy J.C. Anderson, Philip J. Rosenthal, Didier Ménard, David A. Fidock

**Affiliations:** Department of Microbiology and Immunology, Columbia University Irving Medical Center, New York, NY, USA; Department of Microbiology and Immunology, University of Otago, Dunedin, New Zealand; Malaria Genetics and Resistance Unit, INSERM U1201, Institut Pasteur, Paris, France; Institut Cochin Inserm U1016, Université Paris Descartes, Paris, France; Department of Genetics, University of North Carolina at Chapel Hill, Chapel Hill, NC, USA; Programme National de Lutte Contre le Paludisme au Tchad, Ndjamena, Chad; Centre National de Recherche et de Formation sur le Paludisme, Ouagadougou, Burkina Faso; Laboratory of Parasitology, Institute Pasteur of Bangui, Bangui, Central African Republic; University Teaching Hospital of Kamenge, Bujumbura, Burundi; Ifakara Health Institute, Dar es Salaam, Tanzania; Malaria and Other Parasitic Diseases Division, Rwanda Biomedical Centre, Kigali, Rwanda; National Malaria Control Programme, Sierra Leone; National Malaria Control Program, Banjul, The Gambia; Programme National de Lutte Contre le Paludisme, Brazzaville, République du Congo; School of Public Health and Community Medicine, University of Gothenburg, Sweden; National Center for Parasitology, Entomology & Malaria Control, Phnom Penh, Cambodia; Shoklo Malaria Research Unit, Mahidol-Oxford Tropical Medicine Research Unit, Faculty of Tropical Medicine, Mahidol University, Mae Sot, Thailand; Centre for Tropical Medicine and Global Health, Nuffield Department of Medicine, University of Oxford, Oxford, UK; Texas Biomedical Research Institute, San Antonio, TX, USA; Department of Medicine, University of California, San Francisco, CA, USA; Division of Infectious Diseases, Department of Medicine, Columbia University Irving Medical Center, New York, NY, USA

## Abstract

The emergence of artemisinin (ART) resistance in *Plasmodium falciparum* parasites, driven by K13 mutations, has led to widespread antimalarial treatment failure in Southeast Asia. In Africa, our genotyping of 3,299 isolates confirms the emergence of the K13 R561H variant in Rwanda and reveals the continuing dominance of wild-type K13 across 11 countries. We show that this mutation, along with M579I and C580Y, confers varying degrees of *in vitro* ART resistance in African parasites. C580Y and M579I cause substantial fitness costs, which may counter-select against their dissemination in high-transmission settings. We also define the impact of multiple K13 mutations on ART resistance and fitness in multiple Southeast Asian strains. ART susceptibility is unaltered upon editing point mutations in ferrodoxin or mdr2, earlier resistance markers. These data point to the lack of an evident biological barrier to mutant K13 mediating ART resistance in Africa, while identifying their detrimental impact on parasite growth.

## Introduction

Despite recent advances in chemotherapeutics, diagnostics and vector control measures, malaria continues to exert a significant impact on human health. In 2019, cases were estimated at 229 million, resulting in 409,000 fatal outcomes, primarily in Sub-Saharan Africa as a result of *Plasmodium falciparum* infection (WHO, 2020). This situation is predicted to rapidly worsen as a result of the ongoing SARS-CoV-2 pandemic that has crippled malaria treatment and prevention measures (Sherrard-Smith *et al*., 2020). Absent an effective vaccine, malaria control and elimination strategies are critically reliant on the continued clinical efficacy of first-line artemisinin-based combination therapies (ACTs). These ACTs pair fast-acting artemisinin (ART) derivatives with partner drugs such as lumefantrine, amodiaquine, mefloquine or piperaquine (PPQ). ART derivatives can reduce the biomass of drug-sensitive parasites by up to 10,000-fold within 48 h (the duration of one intra-erythrocytic developmental cycle); however, these derivatives are rapidly metabolized *in vivo*. Longer-lasting albeit slower-acting partner drugs are co-administered to reduce the selective pressure for ART resistance and to clear residual parasitemias.

Nonetheless, *P. falciparum* partial resistance to ART derivatives has spread throughout Southeast Asia (SEA), having first emerged a decade ago in western Cambodia (Dondorp *et al*., 2009; Noedl *et al*., 2009; Ariey *et al*., 2014; Imwong *et al*., 2020). Clinically, partial ART resistance manifests as delayed clearance of circulating asexual blood stage parasites following treatment with an ACT. The accepted threshold for resistance is a parasite clearance half-life (the time required for the peripheral blood parasite density to decrease by 50%) of >5.5 h. Sensitive parasites are typically cleared in <2-3 h (WHO, 2019). Partial resistance can also be evidenced as parasite-positive blood smears on day three post initiation of treatment.

*In vitro,* ART resistance manifests as increased survival of cultured ring-stage parasites exposed to a 6 h pulse of 700 nM dihydroartemisinin (DHA, the active metabolite of all ARTs used clinically) in the ring-stage survival assay (RSA) (Witkowski *et al*., 2013; Ariey *et al*., 2014). Recently, ART-resistant strains have also acquired resistance to PPQ, which is widely used in SEA as a partner drug in combination with DHA. Failure rates following DHA-PPQ treatment now exceed 50% in parts of Cambodia, Thailand and Vietnam (van der Pluijm *et al*., 2019).

*In vitro* selections, supported by clinical epidemiological data, have demonstrated that ART resistance is primarily determined by mutations in the beta-propeller domain of the *P. falciparum* Kelch protein K13 (Ariey *et al*., 2014; Ashley *et al*., 2014; MalariaGEN, 2016; Menard *et al*., 2016; Siddiqui *et al*., 2020). Recent evidence suggests that these mutations result in reduced endocytosis of host-derived hemoglobin and thereby decreased release of the ART-activating moiety Fe^2+^-heme, thus reducing ART potency (Yang *et al*., 2019; Birnbaum *et al*., 2020). Mutations in other genes including ferredoxin *(fd)* and multidrug resistance protein 2 *(mdr2)* have also been associated with ART resistance in K13 mutant parasites, suggesting that they either contribute to a multigenic basis of resistance or fitness or serve as genetic markers of founder populations (Miotto *et al*., 2015).

In SEA, the most prevalent K13 mutation is C580Y, which associates with delayed clearance *in vivo* (Ariey *et al*., 2014; Ashley *et al*., 2014; MalariaGEN, 2016; Menard *et al*., 2016; Imwong *et al*., 2017). This mutation also mediates ART resistance *in vitro,* as demonstrated by RSAs on gene-edited parasites (Ghorbal *et al*., 2014; Straimer *et al*., 2015; Uwimana *et al*., 2020). Other studies have since demonstrated the emergence of almost 200 K13 mutations both in SEA and other malaria-endemic regions, including India, the Guiana Shield and the western Pacific (MalariaGEN, 2016; Menard *et al*., 2016; Das *et al*., 2019; Group, 2019; Mathieu *et al*., 2020; Miotto *et al*., 2020). Aside from C580Y, only a handful of K13 mutations (N458Y, M476I, Y493H, R539T, I543T and R561H) have been validated by gene-editing experiments as conferring resistance *in vitro* (Straimer *et al*., 2015; Siddiqui *et al*., 2020). Multiple other mutations have been associated with the clinical delayed clearance phenotype and have been proposed as candidate markers of ART resistance (Group, 2019; WHO, 2019). Here, we define the role of a panel of K13 mutations found in field isolates, and address the key question of whether these mutations can confer resistance in African strains. We include the K13 R561H mutation, earlier associated with delayed parasite clearance in SEA (Ashley *et al*., 2014; Phyo *et al*., 2016), and very recently identified at 7-12% prevalence in certain districts in Rwanda (Uwimana *et al*., 2020; Bergmann *et al*., 2021). This study also enabled us to assess the impact of the parasite genetic background, including ferrodoxin and mdr2 as proposed secondary determinants of resistance, on *in vitro* phenotypes. Our results show that K13 mutations can impart ART resistance across multiple Asian and African strains, at levels that vary widely depending on the mutation and the parasite genetic background. Compared to K13 mutant Asian parasites, we observed stronger *in vitro* fitness costs in K13-edited African strains, which might predict a slower dissemination of ART resistance in high-transmission African settings.

## Results

### Non-synonymous K13 mutations are present at low frequencies in Africa

To examine the status of K13 mutations across Africa, we sequenced the beta-propeller domain of this gene in 3,299 isolates from 11 malaria-endemic African countries, including The Gambia, Sierra Leone, and Burkina Faso in West Africa; Chad, Central African Republic, Republic of the Congo, and Equatorial Guinea in Central Africa; and Burundi, Tanzania, Rwanda, and Somalia in East Africa. Samples were collected between 2011 and 2019, with most countries sampled across multiple years.

Of all samples, 99% (3,220) were K13 WT, i.e. they matched the 3D7 (African) reference sequence or harbored a synonymous (non-coding) mutation. For individual countries, the percentage of K13 WT samples ranged from 95% to 100% (Figure 1; Figure 1–Source data 1). In total, we identified 36 unique non-synonymous mutations in K13. Only two of these non-synonymous mutations have been validated as resistance mediators in the Southeast Asian Dd2 strain: the M476I mutation initially identified from long-term ART selection studies and the R561H mutation observed in Rwanda (Ariey *et al*., 2014; Straimer *et al*., 2015; Uwimana *et al*., 2020).

**Figure 1.**
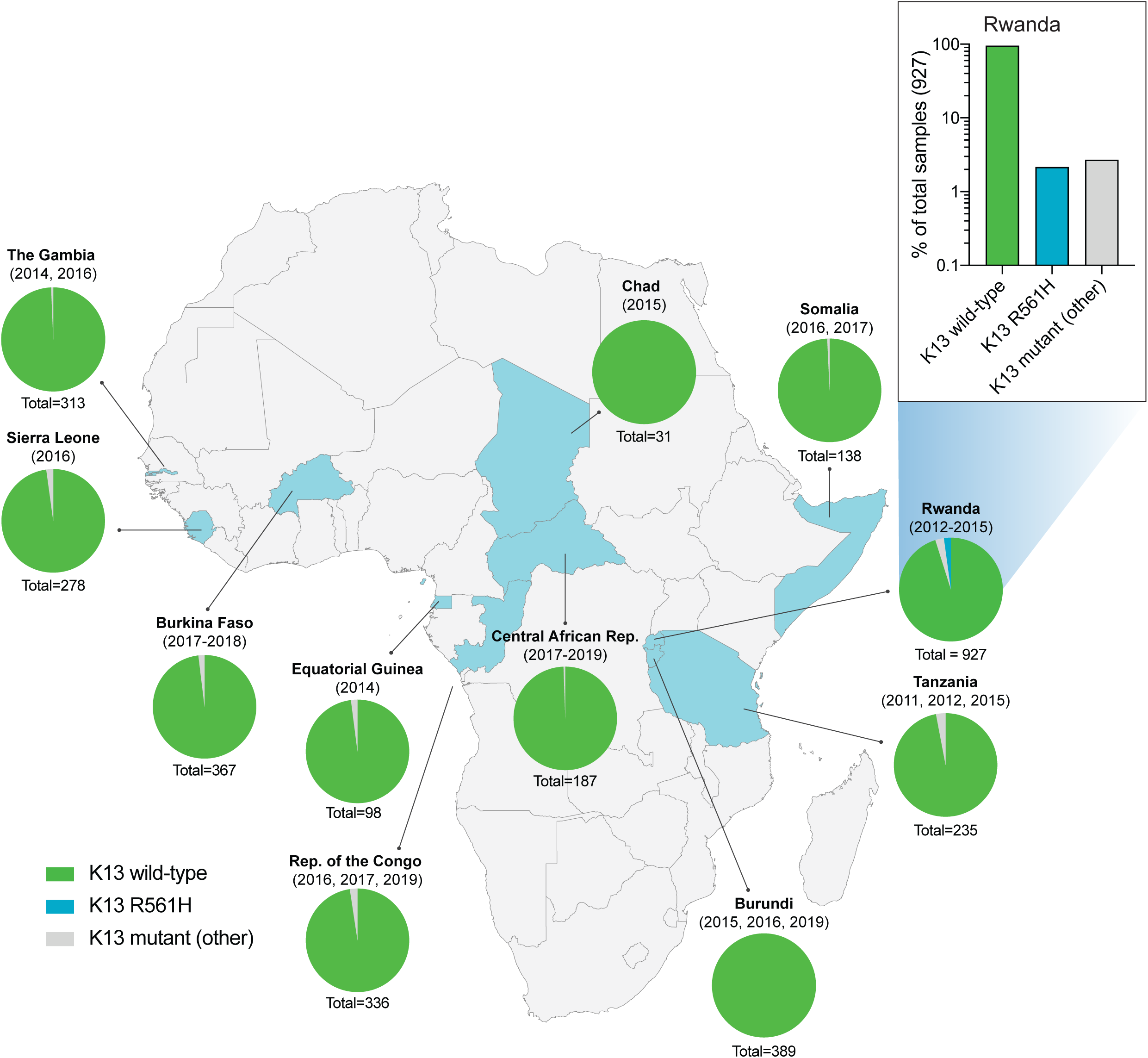
Frequency and distribution of *K13* alleles in 11 African countries. Pie charts representing the proportions of sequenced samples per country that harbor the K13 wild-type (WT; 3D7 reference) sequence, the R561H variant (the most commonly identified mutation, unique to Rwanda; see inset), or other less frequent non-synonymous K13 mutations. Sample sizes and years of sample collection are indicated. Mutations and numbers of African samples sequenced per country are listed in Figure 1–source data 1.

Of the 36 non-synonymous mutations, only two were present in >6 samples: R561H (n=20, unique to Rwanda, sampled from 2012 to 2015), and A578S (n=10; observed in four countries across multiple years). Previously A578S was shown not to confer *in vitro* resistance in Dd2 (Menard *et al*., 2016). R561H accounted for 44% of mutant samples and 7% of all samples in the set of 927 Rwandan genotyped isolates (Uwimana *et al*., 2020).

### The K13 R561H, M579I and C580Y mutations can confer *in vitro* artemisinin resistance in African parasites

To test whether R561H can mediate ART resistance in African strains, we developed a CRISPR/Cas9-mediated *K13* editing strategy (Supplementary file 1) to introduce this mutation into 3D7 and F32 parasites. On the basis of whole-genome sequence analysis of African isolates, 3D7 was recently shown to segregate phylogenetically with parasites from Rwanda (Ariey *et al*., 2014; Uwimana *et al*., 2020). F32 was derived from an isolate from Tanzania (Witkowski *et al*., 2010). We also tested the C580Y mutation that predominates in SEA, as well as the M579I mutation identified in a *P. falciparum*-infected individual in Equatorial Guinea who displayed delayed parasite clearance following ACT treatment (Lu *et al*., 2017). The positions of these residues are highlighted in the K13 beta-propeller domain structure shown in Supplementary file 2. For 3D7, F32 and other lines used herein, the geographic origins and genotypes at drug resistance loci are described in Table 1 and Table 1–table supplement 1. All parental lines were cloned by limiting dilution prior to transfection. Edited parasites were identified by PCR and Sanger sequencing, and cloned.

**Table 1.** Plasmodium falciparum lines employed herein.

| Parasite | Origin | Year | K13 | Resistance |
| --- | --- | --- | --- | --- |
| 3D7<sup>WT</sup> | Africa | 1981 | WT | -- |
| F32<sup>WT</sup> | Tanzania | 1982 | WT | -- |
| UG659<sup>WT</sup> | Uganda | 2007 | WT | CQ, SP |
| UG815<sup>WT</sup> | Uganda | 2008 | WT | CQ, SP |
| Dd2<sup>WT</sup> | Indochina | 1980 | WT | CQ, MFQ, SP |
| Cam3.II<sup>WT</sup> | Cambodia | 2010 | WT | CQ, SP |
| CamWT<sup>C580Y</sup> | Cambodia | 2010 | C580Y | ART, CQ, SP |
| RF7<sup>C580Y</sup> | Cambodia | 2012 | C580Y | ART, CQ, PPQ, SP |
| Thai1<sup>WT</sup> | Thailand | 2003 | WT | CQ, SP |
| Thai2<sup>WT</sup> | Thailand | 2004 | WT | CQ, MFQ, SP |
| Thai3<sup>WT</sup> | Thailand | 2003 | WT | CQ, SP |
| Thai4<sup>WT</sup> | Thailand | 2003 | WT | CQ, SP |
| Thai5<sup>WT</sup> | Thailand | 2011 | WT | CQ, SP |
| Thai6<sup>E252Q</sup> | Thailand | 2008 | E252Q | ART (low), CQ, MFQ, SP |
| Thai7<sup>E252Q</sup> | Thailand | 2010 | E252Q | ART (low), CQ, MFQ, SP |

ART, artemisinin; CQ, chloroquine; MFQ, mefloquine; PPQ, piperaquine; SP, sulfadoxine/pyrimethamine; WT, wild type.

RSAs, used to measure *in vitro* ART susceptibility, revealed a wide range of mean survival values for K13 mutant lines. For 3D7 parasites, the highest RSA survival rates were observed with 3D7^R561H^ parasites, which averaged 6.6% RSA survival. For the 3D7^M579I^ and 3D7^C580Y^ lines, mean RSA survival rates were both 4.8%, a 3 to 4-fold increase relative to the 3D7^WT^ line. No elevated RSA survival was seen in a 3D7 control line (3D7^ctrl^) that expressed only the silent shield mutations used at the guide RNA cut site (Figure 2A; Figure 2–source data 1).

**Figure 2.**
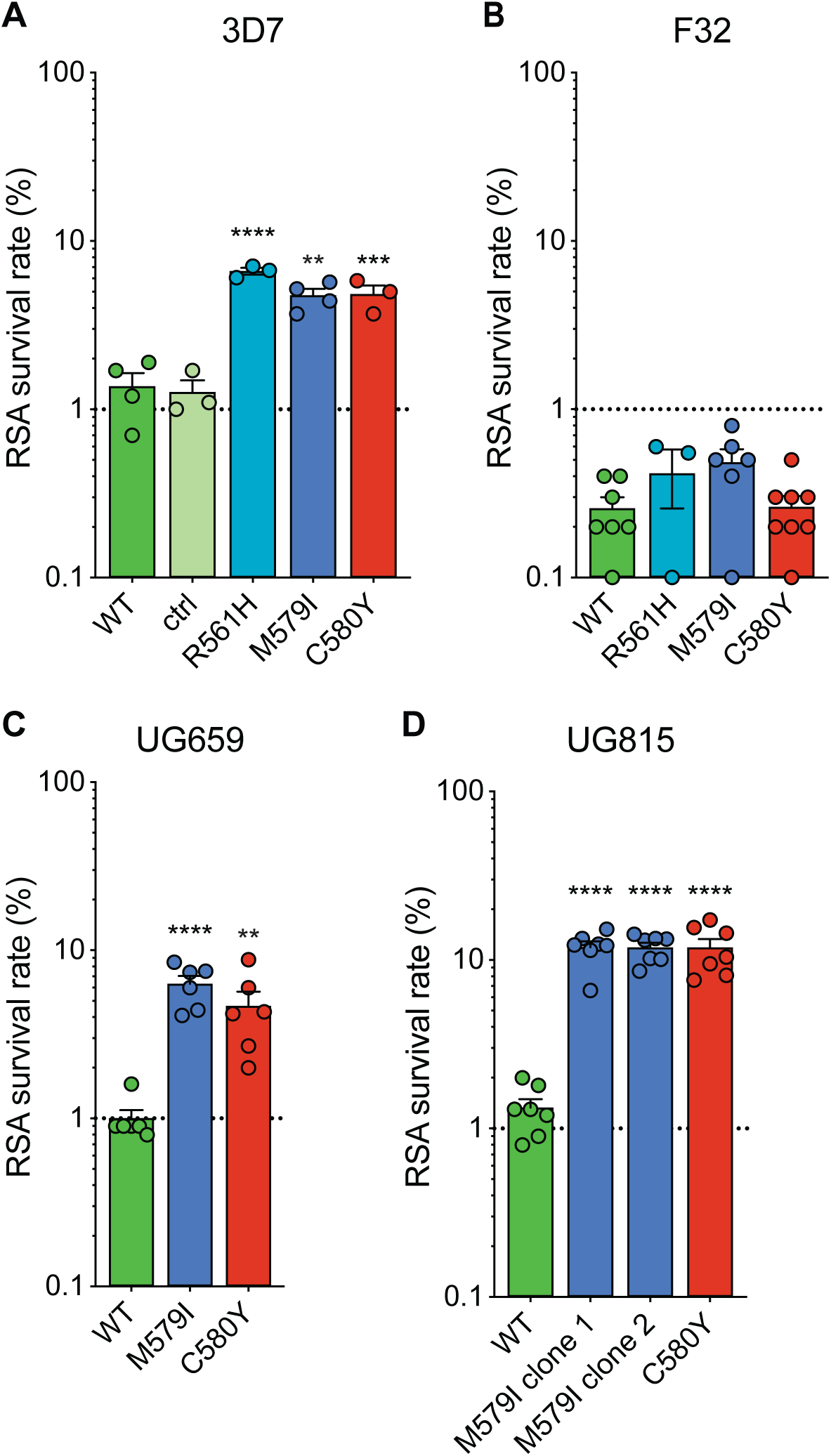
Gene-edited African parasites acquire variable degrees of *in vitro* ART resistance conferred by mutant K13. (A-D) RSA survival rates for (A) 3D7 (Africa), (B) F32 (Tanzania), (C) UG659 (Uganda), or (D) UG815 (Uganda) K13 wild-type parental lines and CRISPR/Cas9-edited K13 R561H, M579I or C580Y mutant clones. Parental lines are described in **Table 1** and Table 1–table supplement 1. For 3D7, we also included a K13 wild-type control (ctrl) line harboring silent-binding site mutations at the K13 gRNA cut site. Results show the percentage of early ring-stage parasites (0-3 hpi) that survived a 6 h pulse of 700 nM DHA, relative to DMSO-treated parasites assayed in parallel. Percent survival values are shown as means ± SEM (detailed in Figure 2–source data 1). Results were obtained from 3 to 8 independent experiments, each performed in duplicate. *P* values were determined by unpaired *t* tests and were calculated for mutant lines relative to the isogenic line expressing WT *K13*. ** *P*<0.01; *** *P*<0.001; **** *P*<0.0001.

In contrast to results with 3D7, the introduction of K13 mutations into F32^WT^ parasites yielded almost no increase in RSA survival. Mean RSA survival rates were 0.5%, 0.5% and 0.3% for F32^R561H^, F32^M579I^ and F32^C580Y^ parasites, respectively, compared to 0.3% for F32^WT^ (Figure 2B). Previously we reported that introduction of M476I into F32 parasites resulted in a modest gain of resistance (mean survival of 1.7%) while this same mutation conferred RSA survival levels of ∼10% in edited Dd2 parasites (Straimer *et al*., 2015). These data suggest that while K13 mutations differ substantially in the level of resistance that they impart, there is an equally notable contribution of the parasite genetic background.

We next introduced M579I and C580Y into cloned Ugandan isolates UG659 and UG815. Editing of both mutations into UG659 yielded moderate RSA survival rates (means of 6.3% and 4.7% for UG659^M579I^ or UG659^C580Y^ respectively, vs. 1.0% for UG659^WT^; Figure 2C). These values resembled our results with 3D7. Strikingly, introducing K13 M579I or C580Y into UG815 yielded the highest rates of *in vitro* resistance, with mean survival levels reaching ∼12% in both UG815^M579I^ and UG815^C580Y^. These results were confirmed in a second independent clone of UG815^M579I^ (Figure 2D). M579I and C580Y also conferred equivalent levels of resistance in edited Dd2 parasites (RSA survival rates of 4.0% and 4.7%, respectively; Figure 2–source data 1). These data show that mutant K13-mediated ART resistance in African parasites can be achieved, in some but not all strains, at levels comparable to or above those seen in Southeast Asian parasites.

### The K13 C580Y and M579I mutations are associated with an *in vitro* fitness defect in African parasites

To examine the relation between resistance and fitness in African parasites harboring K13 mutations, we developed an *in vitro* fitness assay that uses quantitative real-time PCR (qPCR) for allelic discrimination. Assays were conducted with the eight pairs of K13 WT and either C580Y or M579I isogenic parasites used for RSAs, namely 3D7^WT^ + either 3D7^M579I^ or 3D7^C580Y^; F32^WT^ + either F32^M579I^ or F32^C580Y^; UG659^WT^ + either UG659^M579I^ or UG659^C580Y^; and UG815^WT^ + either UG815^M579I^ or UG915^C580Y^.

Assays were initiated with tightly synchronized trophozoites, mixed in 1:1 ratios of WT to mutant isogenic parasites, and cultures were maintained over a period of 40 days (∼20 generations of asexual blood stage growth). Cultures were sampled every four days for genomic DNA (gDNA) preparation and qPCR analysis. TaqMan probes specific to the *K13* WT or mutant (M579I or C580Y) alleles were used to quantify the proportion of each allele.

Results showed that both K13 mutations (M579I or C580Y) conferred a significant fitness defect in all backgrounds tested, with the percentage of the *K13* mutant allele declining over time in all co-cultures (Figure 3; Figure 3–source data 1). This fitness defect varied between parasite backgrounds. To quantify this impact, we calculated the fitness cost, which represents the percent reduction in growth rate per 48 h generation of the test line compared to the WT isogenic comparator. The fitness cost was calculated using day 0 and day 32 as start and end points, respectively, as these yielded the most consistent slopes across lines and time series. For 3D7 parasites, the fitness cost was 8.9% and 6.9% for both the M579I and C580Y mutations, respectively (Figure 3A). In F32 and UG659 parasites, the fitness cost for the M579I mutation was slightly higher (4.3% and 5.8%) than for C580Y (2.8% and 1.9%; Figure 3B, C). The largest growth defects for both mutations were seen in the UG815 line (Figure 3D), with fitness cost values for the M579I and C580Y mutations both being 12.0%. A comparison of data across these four African strains revealed that high RSA survival rates were generally accompanied by high fitness costs, with M579I mostly having a more detrimental fitness impact than C580Y (Figure 3E, 3F).

**Figure 3.**
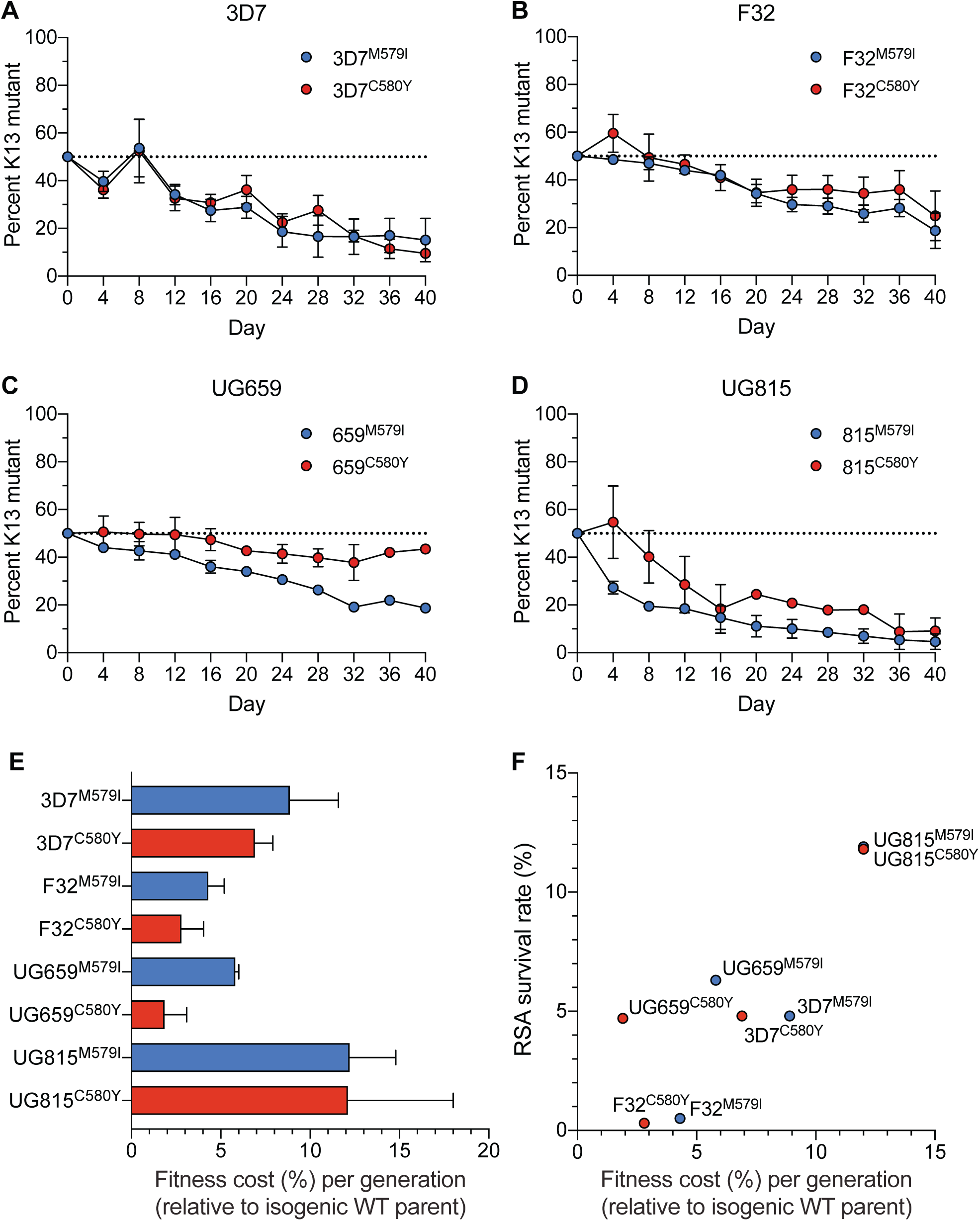
K13 M579I and C580Y mutations cause *in vitro* growth defects in gene-edited African parasites. (A-D) Percentage of mutant allele relative to the WT allele over time in (A) 3D7, (B) F32, (C) UG659, and (D) UG815 parasite cultures in which K13 M579I or C580Y mutant clones were co-cultured as 1:1 mixtures with isogenic K13 WT controls over a period of 40 days. Percentages of mutant and WT alleles in culture over time were determined by TaqMan qPCR-based allelic discrimination, and normalized to 50% on Day 0. Results, shown as means ± SEM, were obtained from 2 to 5 independent experiments, each performed in duplicate. Values are provided in Figure 3–source data 1. (E) The percent reduction in growth rate per 48 h generation, termed the fitness cost, is presented as mean ± SEM for each mutant line relative to its isogenic wild-type comparator. (F) Fitness costs for mutant lines and isogenic wild-type comparators are plotted relative to RSA survival values for the same lines.

### The K13 C580Y mutation has swept rapidly across Cambodia, displacing other K13 variants

We next examined the spatiotemporal distribution of *K13* alleles in Cambodia, the epicenter of ART resistance in SEA. In total, we sequenced the K13 propeller domains of 3,327 parasite isolates collected from western, northern, eastern and southern provinces of Cambodia (Figure 4–figure supplement 1). Samples were collected between 2001 and 2017, except for the southern region where sample collection was initiated in 2010. In sum, 19 nonsynonymous polymorphisms in K13 were identified across all regions and years. Of these, only three were present in >10 samples, Y493H (n=83), R539T (n=87) and C580Y (n=1,915). Each of these mutations was previously shown to confer ART resistance *in vitro* (Straimer *et al*., 2015). Rarer mutations included A418V, I543T, P553L, R561H, P574L, and D584V (Figure 4; Figure 4–source data 1).

**Figure 4.**
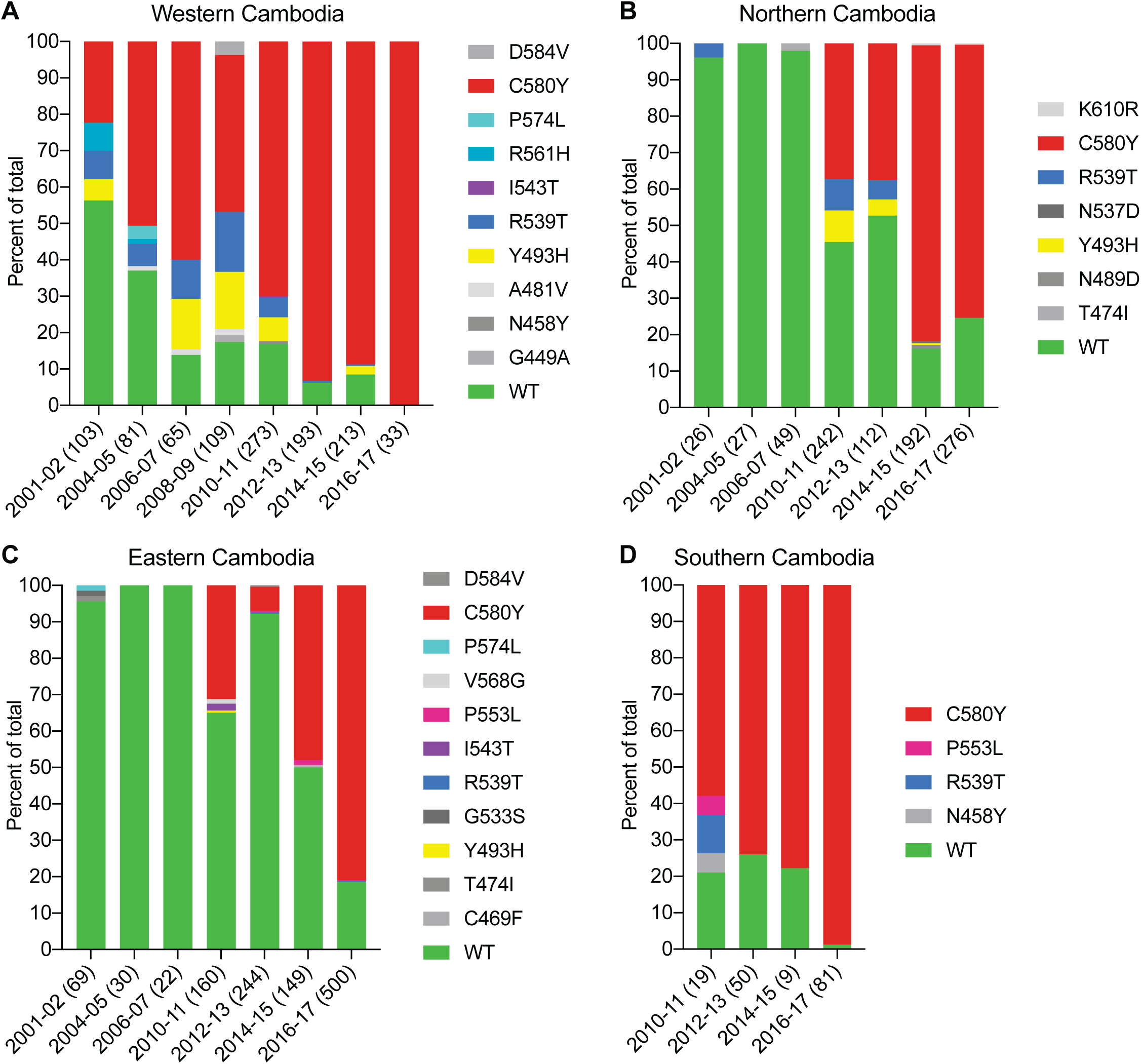
The *K13* C580Y allele has progressively outcompeted all other alleles in Cambodia. (A-D) Stacked bar charts representing the percentage of sequenced samples expressing the *K13* wild-type allele or individual variants, calculated based on the total number of samples (listed in parentheses) for a given period. Sample collection was segregated into four regions in Cambodia (detailed in Figure 4–figure supplement 1). All K13 mutant samples harbored a single non-synonymous nucleotide polymorphism. Mutations and numbers of Cambodian samples sequenced per region/year are listed in Figure 4–source data 1.

This analysis revealed a significant proportion of K13 WT parasites in the early 2000s, particularly in northern and eastern Cambodia, where 96% of isolates in 2001-2002 had the WT *K13* sequence (Figure 4). In western Cambodia, where ART resistance first emerged (Dondorp *et al*., 2009; Noedl *et al*., 2009), the WT allele percentage in 2001-2002 had already fallen to 56%. This is striking given that delayed parasite clearance following ACT or artesunate treatment was first documented in 2008-2009 (Noedl *et al*., 2008; Noedl *et al*., 2009).

In all four regions, the frequency of the WT allele declined substantially over time and the diversity of mutant alleles contracted, with nearly all WT and non-K13 C580Y mutant parasites being replaced by parasites harboring the C580Y mutation (Figure 4). This effect was particularly pronounced in the west and the south, where the prevalence of C580Y in 2016-17 effectively attained 100%, increasing from 22% and 58% respectively in the initial sample sets (Figure 4A, D). In northern and eastern Cambodia, C580Y also outcompeted all other mutant alleles; however, 19-25% of parasites remained K13 WT in 2016-17 (Figure 4B, C). These data show the very rapid dissemination of K13 C580Y across Cambodia.

### Southeast Asian K13 mutations associated with delayed parasite clearance differ substantially in their ability to confer artemisinin resistance *in vitro*

Given that most K13 polymorphisms present in the field have yet to be characterized *in vitro*, we selected a panel of mutations to test by gene editing, namely E252Q, F446I, P553L, R561H and P574L (Supplementary file 2). The F446I mutation is the predominant mutation in the Myanmar-China border region. P553L, R561H and P574L have each been shown to have multiple independent origins throughout SEA (Menard *et al*., 2016), and were identified at low frequencies in our sequencing study in Cambodia (Figure 4). Lastly, the E252Q mutation was formerly prevalent on the Thai-Myanmar border, and, despite its occurrence upstream of the beta-propeller domain, has been associated with delayed parasite clearance *in vivo* (Anderson *et al*., 2017; Cerqueira *et al*., 2017; Group, 2019).

Zinc-finger nuclease- or CRISPR/Cas9-based gene edited lines expressing K13 E252Q, F4461, P553L, R561H or P574L were generated in Dd2 or Cam3.II lines expressing WT K13 (Dd2^WT^ or Cam3.II^WT^) and recombinant parasites were cloned. Early ring-stage parasites were then assayed for their ART susceptibility using the RSA. For comparison, we included published Dd2 and Cam3.II lines expressing either K13 C580Y (Dd2^C580Y^ and Cam3.II^C580Y^) or R539T (Dd2^R539T^ and the original parental line Cam3.II^R539T^) (Straimer *et al*., 2015), as well as control lines expressing only the guide-specific silent shield mutations (Dd2^ctrl^ and Cam3.II^ctrl^).

Both the P553L and R561H mutations yielded mean RSA survival rates comparable to C580Y (4.6% or 4.3% RSA survival for Dd2^P553L^ or Dd2^R561H^, respectively, vs 4.7% for Dd2^C580Y^; Figure 5A; Figure 5–source data 1). F446I and P574L showed only modest increases in survival relative to the WT parental line (2.0% and 2.1% for Dd2^F446I^ and Dd2^P574L^, respectively, vs 0.6% for Dd2^WT^). No change in RSA survival relative to Dd2^WT^ was observed for the Dd2^E252Q^ line. The resistant benchmark Dd2^R539T^ showed a mean RSA survival level of 20.0%.

**Figure 5.**
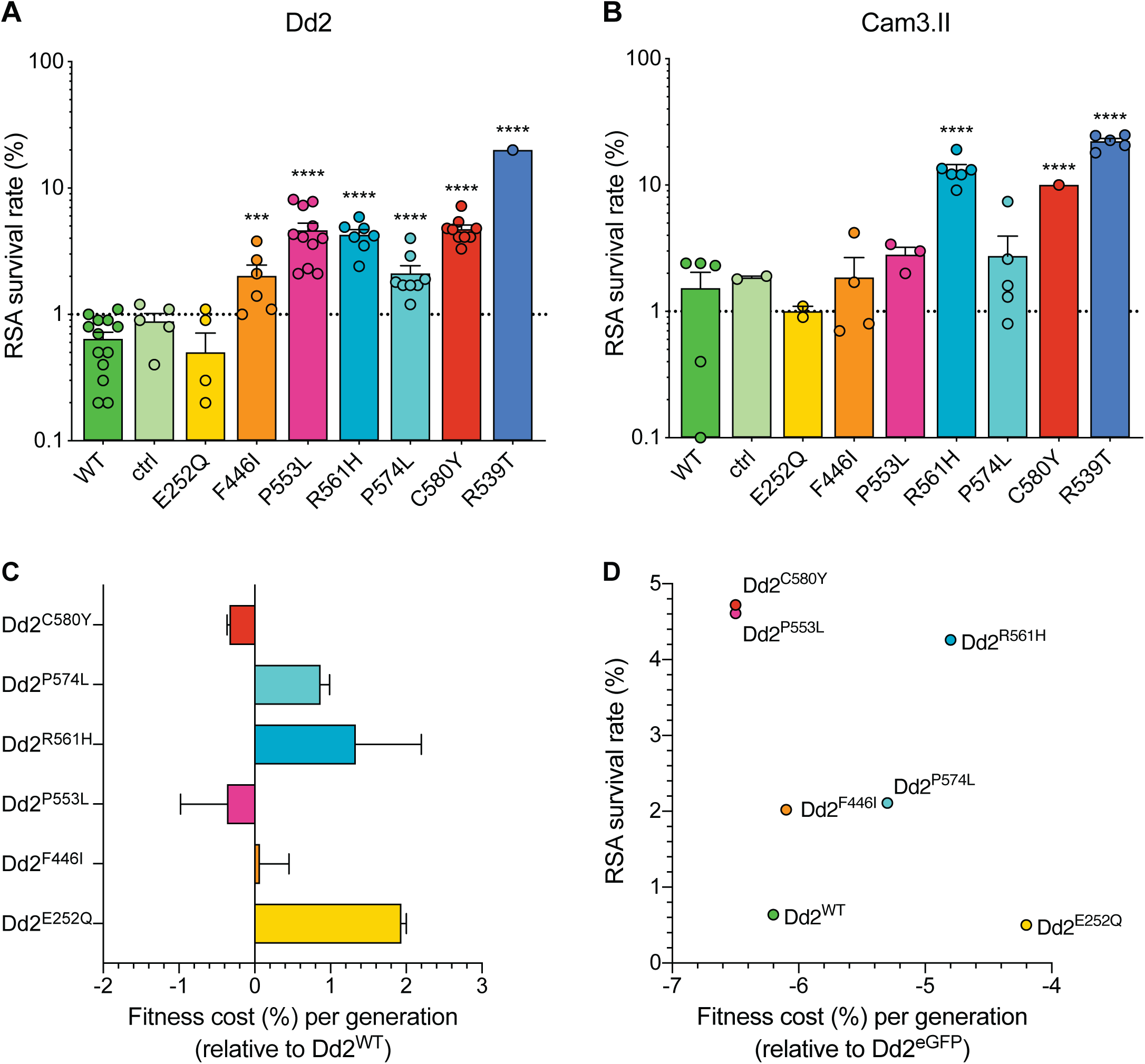
Southeast Asian K13 mutations yield elevated RSA survival and minor impacts on *in vitro* growth in gene-edited parasite lines. (A, B) RSA survival rates for Dd2 (Indochina) and Cam3.II (Cambodia) *P. falciparum* parasites expressing wild-type or mutant K13. Gene-edited parasites were generated using CRISPR/Cas9 or zinc-finger nucleases (ZFNs). Control (ctrl) lines express silent-binding site mutations at the *K13* gRNA cut site. Unedited parental lines are described in **Table 1** and Table 1–table supplement 1. Results show the percentage of early ring-stage parasites (0-3 hpi) that survived a 6 h pulse of 700 nM DHA, relative to DMSO-treated parasites processed in parallel. Percent survival values are shown as means ± SEM (detailed in Figure 5–source data 1). Results were obtained from 3 to 13 independent experiments, each performed in duplicate. *P* values were determined by unpaired *t* tests and were calculated for mutant lines relative to the isogenic line expressing WT *K13*. *** *P*<0.001; **** *P*<0.0001. (C) Percent reductions in growth rate per 48 h generation, expressed as the fitness cost, for each Dd2 mutant line relative to the Dd2^WT^ line. Fitness costs were determined by co-culturing a Dd2^eGFP^ reporter line with either the Dd2 K13 wild-type parental line (Dd2^WT^) or gene-edited K13 mutant lines. Co-cultures were maintained for 20 days and percentages of eGFP^+^ parasites were determined by flow cytometry (see Figure 5–source data 2 and Figure 5–figure supplement 1). Fitness costs were calculated relative to the Dd2^eGFP^ reporter line (Figure 5–figure supplement 1) and are shown here normalized to the Dd2^WT^ line. Mean ± SEM values were obtained from three independent experiments, each performed in triplicate. (D) Fitness costs for K13 WT or mutant lines, relative to the Dd2^eGFP^ line, plotted against their corresponding RSA survival values.

In contrast to Dd2, editing of the F446I, P553L and P574L mutations into the Cambodian Cam3.II parasite background did not result in a statistically significant increase in survival relative to K13 WT Cam3.II^WT^ parasites, in part because the background survival rate of the Cam3.II^WT^ line was higher than for Dd2^WT^. All survival values were <3%, contrasting with the Cam3.II^R539T^ parental line that expresses the R539T mutation (∼20% mean survival; Figure 5B; Figure 5–source data 1). The E252Q mutation did not result in elevated RSA survival in the Cam3.II background, a result also observed with Dd2. Nonetheless, ART resistance was apparent upon introducing the R561H mutation into Cam3.II parasites, whose mean survival rates exceeded the Cam3.II^C580Y^ line (13.2% vs 10.0%, respectively). No elevated survival was seen in the Cam3.II^ctrl^ line expressing only the silent shield mutations used at the guide RNA cut site.

### Southeast Asian K13 mutations do not impart a significant fitness impact on Dd2 parasites

Prior studies with isogenic gene-edited Southeast Asian lines have shown that certain K13 mutations can exert fitness costs, as demonstrated by reduced intra-erythrocytic asexual blood stage parasite growth (Straimer *et al*., 2017; Nair *et al*., 2018). To determine the fitness impact of the K13 mutations described above, we utilized an eGFP-based parasite competitive growth assay (Ross *et al*., 2018). Dd2^E252Q^, Dd2^F446I^, Dd2^P553L^, Dd2^R561H^ or Dd2^P574L^ were co-cultured in 1:1 mixtures with an isogenic K13 WT eGFP^+^ Dd2 reporter line for 20 days (10 generations), and the proportion of eGFP^+^ parasites was assessed every two days. As controls, we included Dd2^WT^, Dd2^bsm^ and Dd2^C580Y^. These data provided evidence of a minimal impact with the F446I, P553L and C580Y mutations, with E252Q, R561H and P574L having greater fitness costs (Figure 5C; Figure 5–figure supplement 1; Figure 5– source data 2). Both C580Y and P553L displayed elevated RSA survival and minimal fitness cost in the Dd2 strain, providing optimal traits for dissemination (Figure 5D). We note that all fitness costs in Dd2 were considerably lower than those observed in our four African strains (Figure 3).

### Strain-dependent genetic background differences significantly RSA survival rates in culture-adapted Thai isolates

Given the earlier abundance of the R561H and E252Q alleles in border regions of Thailand and Myanmar, we next tested the impact of introducing these mutations into five Thai K13 WT isolates (Thai1-5). For comparison, we also edited C580Y into several isolates. These studies revealed a major contribution of the parasite genetic background in dictating the level of mutant K13-mediated ART resistance, as exemplified by the C580Y lines whose mean survival rates ranged from 2.1% to 15.4%. Trends observed for individual mutations were maintained across strains, with the R561H mutation consistently yielding moderate to high levels of *in vitro* resistance at or above the level of C580Y. Consistent with results for Dd2, introduction of E252Q did not result in significant increases in survival rates relative to isogenic K13 WT lines (Figure 6A-E; Figure 6–source data 1).

**Figure 6.**
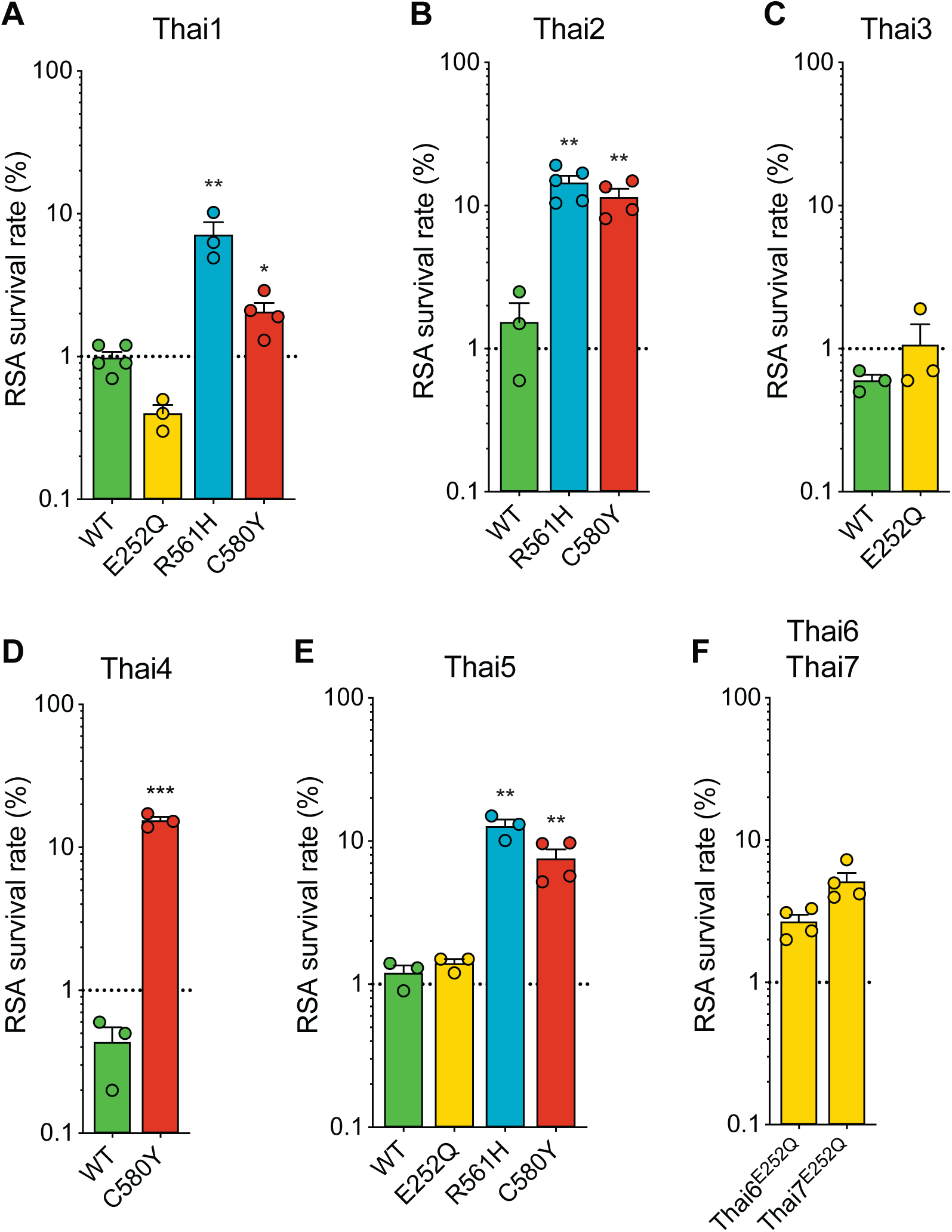
Thai isolates expressing mutant K13 display variable RSA survival rates. RSA survival rates for (A-E) *K13*-edited Thai isolates and (F) K13 E252Q unedited Thai lines, shown as means ± SEM (detailed in Figure 6–source data 1). Results were obtained from 3 to 7 independent experiments, each performed in duplicate. *P* values were determined by unpaired *t* tests and were calculated for mutant lines relative to the isogenic line expressing WT *K13*. * *P*<0.05; ** *P*<0.01; *** *P*<0.001.

We also profiled two unedited culture-adapted Thai isolates (Thai6^E252Q^ and Thai7^E252Q^) that express the K13 E252Q mutation, but that are otherwise K13 WT. Notably, both lines exhibited mean RSA survival rates significantly above the 1% threshold for ART sensitivity (2.7% for Thai6^E252Q^ and 5.1% for Thai7^E252Q^; Figure 6F). These data suggest that additional genetic factors present in these two Thai isolates are required for E252Q to manifest ART resistance.

### Mutations in the *P. falciparum* multidrug resistance protein 2 and ferredoxin genes do not modulate resistance to artemisinin *in vitro*

In a prior genome-wide association study of SE Asian parasites, K13-mediated ART resistance was associated with D193Y and T484I mutations in the ferredoxin (*fd*) and multidrug resistance protein 2 (*mdr2*) genes, respectively (Miotto *et al*., 2015). To directly test the role of these mutations, we applied CRISPR/Cas9 editing to the Cambodian strains RF7^C580Y^ and Cam3.II^C580Y^, which both express K13 C580Y (Supplementary file 3). Isogenic RF7 parasites expressing the mutant or wild-type fd residue at position 193 showed no change in RSA survival rates, either at 700 nM (averaging ∼27%), or across a range of DHA concentrations down to 1.4 nM. Editing fd D193Y into K13 C580Y CamWT parasites (Straimer *et al*., 2015) also had no impact on RSA survival (with mean RSA survival rates of 11-13%). Likewise, Cam3.II parasites maintained the same rate *of in vitro* RSA survival (mean 19-22%) irrespective of their *mdr2* allele. Silent shield mutations had no impact for either *fd* or *mdr2* (Figure 7; Figure 7–source data 1). These data suggest that the fd D193Y and mdr2 T484I mutations are markers of ART-resistant founder populations but themselves do not contribute directly to ART resistance.

**Figure 7.**
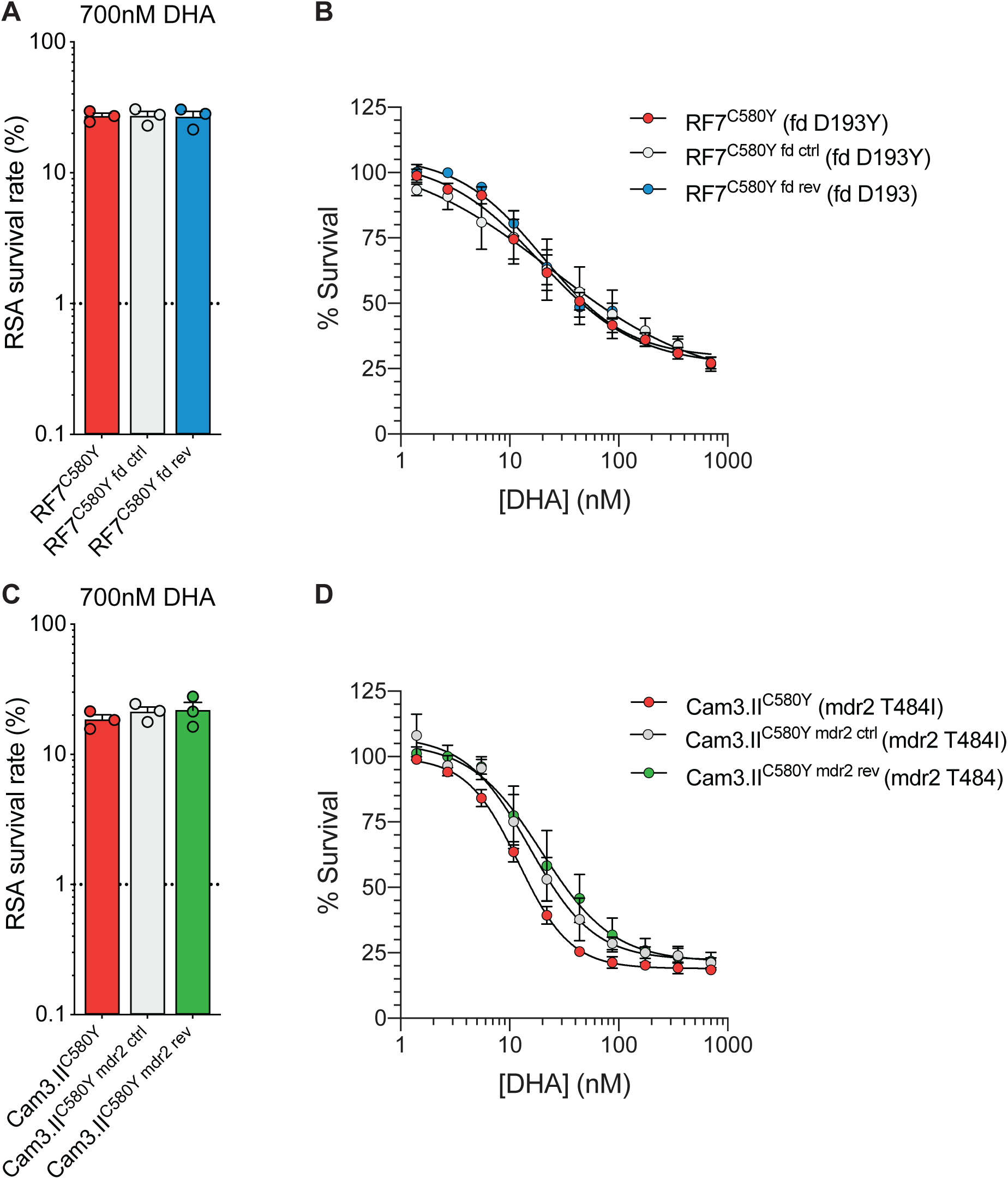
Ferrodoxin (*fd)* and multidrug resistance protein 2 (*mdr2)* mutations do not impact RSA survival in K13 C580Y parasites. (A, B) Results of RSA assays performed on the RF7^C580Y^ parental line that expresses the fd variant D193Y, a gene-edited control line with silent binding-site mutations at the *fd* gRNA cut site, and a gene-edited fd revertant D193 line. Gene-edited parasite lines were generated using CRISPR/Cas9. Early ring-stage parasites (0-3 hpi) were pulsed with serial dilutions of DHA (beginning with 700 nM DHA) for 6 h followed by drug washout. Results show (A) mean ± SEM survival at 700 nM, and (B) survival curves across the range of DHA concentrations. Survival rates were calculated relative to DMSO-treated parasites processed in parallel. Results were obtained from three independent experiments, each performed in duplicate. (C, D) RSAs performed on the Cam3.II^C580Y^ parental line (expressing the mdr2 variant T484I), a gene-edited control line with silent-binding site mutations at the *mdr2* gRNA cut site, and a gene-edited mdr2 revertant T484 line. Results shown in (C, D) were generated and presented as per (A, B). All values are provided in Figure 7–source data 1.

## Discussion

Mutant K13-mediated ART resistance has substantially compromised the efficacy of antimalarial treatments across SEA, and the relatively high prevalence of the R561H variant associated with delayed clearance in Rwanda now poses a threat to high-transmission settings in sub-Saharan Africa (Conrad and Rosenthal, 2019; Hanboonkunupakarn and White, 2020; Uwimana *et al*., 2020; Bergmann *et al*., 2021). Using gene editing and phenotypic analyses, we provide definitive evidence that the K13 R561H, M579I and C580Y mutations can confer *in vitro* ART resistance in several African strains. *In vitro* resistance, as defined using the RSA, was comparable between gene-edited African K13 R561H 3D7 parasites and Asian C580Y Dd2 and Cam3.II parasites. We also observed that K13 mutant African strains differed widely in their RSA survival rates. As an example, when introduced into the Tanzanian F32 and Ugandan UG815 strains, the C580Y mutation yielded 0.3% (not resistant) and 11.8% (highly resistant) RSA survival rates, respectively. These data suggest that F32 parasites lack additional genetic determinants that are required for mutant K13 to confer ART resistance. Nonetheless, our results provide conclusive evidence that multiple African strains present no core biological obstacle to becoming ART resistant upon acquiring K13 mutations.

Our spatio-temporal analysis of K13 sequence diversity in Cambodia highlights the emergence of C580Y in western Cambodia and its progressive replacement of other variants (Imwong *et al*., 2020). The success of this mutation in SEA cannot be explained by resistance alone, as we previously reported that the less common R539T and I543T variants conferred greater ART resistance *in vitro* (Straimer *et al*., 2015). Similarly, we now report that R561H and P553L yield equivalent degrees of ART resistance when compared with C580Y. Lower levels of resistance were observed with F446I and P574L, with the former predominating recently on the Thai-Myanmar border (Imwong *et al*., 2020). In a recent study, F446I yielded no significant *in vitro* resistance in 3D7 parasites, although similar to our data this mutation was fitness-neutral (Siddiqui *et al*., 2020). Of note, all four of these mutations, and others including C469Y, R622I and A675V, have now been detected in Africa and merit gene editing experiments in African strains (Warsame *et al*., 2019; Asua *et al*., 2020; Kayiba *et al*., 2020).

Here we observed that the C580Y mutation exerts less of a fitness cost relative to other K13 variants, as measured in *K13*-edited Dd2 parasites co-cultured with an eGFP reporter line. These data suggest that C580Y might be favored in part because of a less detrimental impact on asexual blood stage growth rates. The most detrimental impact on growth was observed with E252Q, which earlier predominated near the Thailand-Myanmar border but was later overtaken by C580Y, as well as R561H, which progressively disappeared over time in SEA (Phyo *et al*., 2016). In our study C580Y produced an optimal combination of no measurable fitness cost and relatively high RSA survival rates in Dd2 parasites. R561H, however, showed slightly improved fitness relative to C580Y in paired isogenic parasites from Thailand (the NHP4302 strain) (Nair *et al*., 2018), providing evidence that both fitness and resistance are strain-dependent.

Consistent with these findings, we observed substantial fitness costs with the K13 C580Y mutation in four African strains. The largest growth defect was observed with the edited UG815 C580Y line that also yielded the highest level of ART resistance. These data suggest that K13 C580Y may not easily take hold in Africa where, unlike in SEA, infections are often highly polyclonal, generating intra-host competition that impacts a strain’s ability to succeed at the population level. In addition, individuals in highly-endemic African settings generally have high levels of acquired immunity, potentially preventing infection by relatively unfit parasites, and often have asymptomatic infections that go untreated and are thus less subject to selective drug pressure, compared with individuals in SEA. This situation recalls the history of chloroquine use in Africa, where fitness costs caused by mutations in the primary resistance determinant PfCRT resulted in the rapid resurgence of wild-type parasites following the implementation of other first-line antimalarial therapies (Kublin *et al*., 2003; Laufer *et al*., 2006; Ord *et al*., 2007; Frosch *et al*., 2014). It remains to be determined whether mutations such as R561H, emerging in Rwanda, can ameliorate the fitness cost observed with other K13 variants in African strains.

Further research is also required to define secondary genetic determinants that could augment mutant K13-mediated ART resistance and to explore other potential mediators of resistance. The latter include mutations in AP-2μ, UBP-1 and Pfcoronin, which can modulate *P. falciparum* ART susceptibility *in vitro* and merit further investigation (Demas *et al*., 2018; Henrici *et al*., 2019; Sutherland *et al*., 2020). Data provided herein argue against a direct role for mutations in *fd* and *mdr2*, earlier associated with mutant K13-mediated resistance in SEA (Miotto *et al*., 2015). We note that *P. falciparum* population structures in Africa tend to be far more diverse than in the epicenter of resistance in Cambodia, where parasite strains are highly sub-structured into a few lineages that can readily maintain complex genetic traits (Amato *et al*., 2018). A requirement to transmit mutant K13 and additional determinants of resistance in African malaria-endemic settings, where genetic outcrossing is the norm, would predict that ART resistance will spread more gradually than in SEA.

Another impediment to the dissemination of ART resistance in Africa is the continued potent activity of lumefantrine, the partner drug in the first line treatment artemether-lumefantrine. This situation contrasts with SEA where ART-resistant parasites also developed high-level resistance to the partner drug PPQ, with widespread treatment failures enabling the dissemination of multidrug-resistant strains (Conrad and Rosenthal, 2019; van der Pluijm *et al*., 2019). These data call for continent-wide monitoring for the emergence and spread of mutant K13 in Africa, and for studies of whether its emergence in Rwanda is a harbinger of subsequent partner drug resistance and ACT treatment failure.

## Materials and Methods

### Sample collection and k13 genotyping

Samples were obtained as blood-spot filter papers from patients seeking treatment at sites involved in national surveys of antimalarial drug resistance, from patients enrolled in therapeutic efficacy studies, from asymptomatic participants who were enrolled in surveillance programs. Collection details from African and Cambodian samples are provided in Figure 1– source data 1 and Figure 4–source data 1, respectively. Samples were processed at the Pasteur Institute in Paris (France) or the Pasteur Institute in Cambodia (Cambodia), as detailed in Supplementary file 4. These investigators vouch for the accuracy and completeness of the molecular data. DNA was extracted from dried blood spots using QIAmp Mini kits, as described (Menard *et al*., 2016). A nested PCR was performed on each sample to amplify the K13-propeller domain, corresponding to codons 440-680. PCR products were sequenced using internal primers and electropherograms analyzed on both strands, using the Pf3D7_1343700 3D7 sequence from 3D7 parasites as the reference sequence. Quality controls included adding six blinded quality-control samples to each 96-well sequencing plate prepared from samples from each in-country partner and independently retesting randomly selected blood samples. Isolates with mixed alleles were considered to be mutated for the purposes of estimating the mutation frequencies.

### *P. falciparum* parasite *in vitro* culture

*Plasmodium falciparum* asexual blood-stage parasites were cultured in human erythrocytes at 3% hematocrit in RPMI-1640 medium supplemented with 2 mM L-glutamine, 50 mg/L hypoxanthine, 25 mM HEPES, 0.225% NaHCO3, 10 mg/L gentamycin and 0.5% w/v Albumax II (Invitrogen). Parasites were maintained at 37°C in 5% O_2_, 5% CO_2_, and 90% N_2_. Cultures were monitored by light microscopy of methanol-fixed, Giemsa-stained blood smears. The geographic origin and year of culture adaptation for lines employed herein are described in **Table 1** and Table 1–table supplement 1.

### Whole-genome sequencing of parental lines

To define the genome sequences of our *P. falciparum* lines used for transfection, we lysed parasites in 0.05% saponin, washed them with 1×PBS, and purified genomic DNA (gDNA) using the QIAamp DNA Blood Midi Kit (Qiagen). DNA concentrations were quantified by NanoDrop (Thermo Scientific) and Qubit (Invitrogen) prior to sequencing. 200 ng of gDNA was used to prepare sequencing libraries using the Illumina Nextera DNA Flex library prep kit with dual indices. Samples were multiplexed and sequenced on an Illumina MiSeq to obtain 300 bp paired-end reads at an average of 50× depth of coverage. Sequence reads were aligned to the *P. falciparum* 3D7 reference genome (PlasmoDB version 36) using Burrow-Wheeler Alignment. PCR duplicates and unmapped reads were filtered out using Samtools and Picard. Reads were realigned around indels using GATK RealignerTargetCreator and base quality scores were recalibrated using GATK BaseRecalibrator. GATK HaplotypeCaller (version 3.8) was used to identify all single nucleotide polymorphisms (SNPs). These SNPs were filtered based on quality scores (variant quality as function of depth QD > 1.5, mapping quality > 40, min base quality score > 18) and read depth (> 5) to obtain high-quality SNPs, which were annotated using snpEFF. Integrated Genome Viewer was used to visually verify the presence of SNPs. BIC-Seq was used to check for copy number variations using the Bayesian statistical model (Xi *et al*., 2011). Copy number variations in highly polymorphic surface antigens and multi-gene families were removed as these are prone to copy number changes with *in vitro* culture.

These whole-genome sequencing data were used to determine the genotypes of the antimalarial drug resistance loci *pfcrt*, *mdr1*, *dhfr* and *dhps* (Haldar *et al*., 2018). We also genotyped *fd*, *arps10*, *mdr2*, *ubp1,* and *ap-2μ*, which were previously associated with ART resistance (Henriques *et al*., 2014; Miotto *et al*., 2015; Cerqueira *et al*., 2017; Adams *et al*., 2018). These results are described in Table 1–table supplement 1.

### Cloning of *K13, fd* and *mdr2* plasmids

Zinc-finger nuclease-meditated editing of select mutations in the *K13* locus was performed as previously described (Straimer *et al*., 2015). CRISPR/Cas9 editing of *K13* mutations was achieved using the pDC2-cam-coSpCas9-U6-gRNA-h*dhfr* all-in-one plasmid that contains a *P. falciparum* codon-optimized Cas9 sequence, a human dihydrofolate reductase (h*dhfr*) gene expression cassette (conferring resistance to WR99210) and restriction enzyme insertion sites for the guide RNA (gRNA) and donor template (White *et al*., 2019). A K13 propeller domain-specific guide gRNA was introduced into this vector at the BbsI restriction sites using the oligo pair p1+p2 (Supplementary file 5) using T4 DNA ligase (New England BioLabs). Oligos were phosphorylated and annealed prior to cloning. A *K13* donor template consisting of a 1.5 kb region of the *K13* coding region including the entire propeller domain was amplified using the primer pair p3+p4 and cloned into the pGEM T-easy vector system (Promega). This donor sequence was subjected to site-directed mutagenesis in the pGEM vector to introduce silent binding-site mutations at the Cas9 cleavage site using the primer pair p5+p6, and to introduce allele-specific mutations using the primer pairs (p7 to p20). *K13* donor sequences were amplified from the pGEM vector using the primer pair p21+p22 and sub-cloned into the pDC2-cam-coSpCas9-U6-gRNA-h*dhfr* plasmid at the EcoRI and AatII restriction sites by In-Fusion® Cloning (Takara). The final plasmids were then sequenced using primers p23 to p25. A schematic showing the method of *K13* plasmid construction can be found in Supplementary file 1.

CRISPR/Cas9 editing of *fd* and *mdr2* was performed using a separate all-in-one plasmid, pDC2-cam-Cas9-U6-gRNA-h*dhfr*, generated prior to the development of the codon-optimized version used above for *K13* (Lim *et al*., 2016). Cloning was performed as for *K13*, except for gRNA cloning that was performed using In-Fusion® cloning (Takara) rather than T4 ligase. Cloning of gRNAs was performed using primer pair p29/p30 for *fd* and p42/p43 for *mdr2*. Donor templates were amplified and cloned into the final vector using the primer pairs p31/p32 for *fd* and p44+p45 for *mdr2*. Site-directed mutagenesis was performed using the allele-specific primer pairs p33+p34 or p35+p36 for *fd*, and p46+p47 or p48+p49 for *mdr2*. All final plasmids (both *fd*- and *mdr2*-specific) were sequenced using the primer pair p37+p38 (Supplementary file 5; Supplementary file 6). Schematic representations of final plasmids are shown in Supplementary file 3.

### Generation of *K13*, *fd* and *mdr2* gene-edited parasite lines

Gene-edited lines were generated by electroporating ring-stage parasites at 5-10% parasitemia with 50 μg of purified circular plasmid DNA resuspended in Cytomix. Transfected parasites were selected by culturing in the presence of WR99210 (Jacobus Pharmaceuticals) for six days post electroporation. Parental lines harboring 2-3 mutations in the *P. falciparum* dihydrofolate reductase (*dhfr*) gene were exposed to 2.5 nM WR99210, while parasites harboring four *dhfr* mutations were selected under 10 nM WR99210 (see Table 1–table supplement 1). Parasite cultures were monitored for recrudescence by microscopy for up six weeks post electroporation. To test for successful editing, the *K13* locus was amplified directly from whole blood using the primer pair p26+p27 (Supplementary file 5) and the MyTaq™ Blood-PCR Kit (Bioline). Primer pairs p39+p40 and p50+p51 were used to amplify *fd* and *mdr2*, respectively. PCR products were submitted for Sanger sequencing using the PCR primers as well as primer p28 in the case of *K13*, p41 (*fd*) or p52 (*mdr2*). Bulk-transfected cultures showing evidence of editing by Sanger sequencing were cloned by limiting dilution.

### Parasite synchronization, ring-stage survival assays (RSAs) and flow cytometry

Synchronized parasite cultures were obtained by exposing predominantly ring-stage cultures to 5% D-Sorbitol (Sigma) for 15 min at 37°C to remove mature parasites. After 36 h of subsequent culture, multinucleated schizonts were either purified over a density gradient consisting of 75% Percoll (Sigma). Purified schizonts were incubated with fresh RBCs for 3h, and early rings (0-3 hours post invasion; hpi) were treated with 5% D-Sorbitol to remove remaining schizonts.

*In vitro* RSAs were conducted as previously described, with minor adaptations (Straimer *et al*., 2015). Briefly, tightly synchronized 0-3 hpi rings were exposed to a pharmacologically-relevant dose of 700 nM DHA or 0.1% dimethyl sulfoxide (DMSO; vehicle control) for 6 h at 1% parasitemia and 2% hematocrit, washed three times with RPMI medium to remove drug, transferred, and cultured for an additional 66 h in drug-free medium. Removal of media and resuspension of parasite cultures was performed on a Freedom Evo 100 liquid-handling instrument (Tecan). Parasitemias were measured at 72 h by flow cytometry (see below) with at least 50,000 events captured per sample. Parasite survival was expressed as the percentage value of the parasitemia in DHA-treated samples divided by the parasitemia in DMSO-treated samples processed in parallel. We considered any RSA mean survival rates <2% to be ART sensitive.

Flow cytometry was performed on an BD Accuri^TM^ C6 Plus cytometer with a HyperCyt plate sampling attachment (IntelliCyt), or on an iQue3® Screener Plus cytometer (Sartorius). Cells were stained with 1×SYBR Green (Invitrogen) and 100 nM MitoTracker DeepRed (Invitrogen) for 30 min and diluted in 1×PBS prior to sampling. Percent parasitemia was determined as the percentage of MitoTracker^-^positive and SYBR Green-positive cells. For RSAs, >50,000 events were captured per well.

### TaqMan allelic discrimination real-time (quantitative) PCR-based fitness assays

Fitness assays with African *K13*-edited parasite lines were performed by co-culturing isogenic wild-type unedited and mutant edited parasites in 1:1 ratios. Assays were initiated with tightly synchronized trophozoites. Final culture volumes were 3 mL. Cultures were maintained in 12-well plates and monitored every four days over a period of 40 days (20 generations) by harvesting at each time point a fraction of each co-culture for saponin lysis. gDNA was then extracted using the QIAamp DNA Blood Mini Kit (Qiagen). The percentage of the WT or mutant allele in each sample was determined in TaqMan allelic discrimination real-time PCR assays. TaqMan primers (forward and reverse) and TaqMan fluorescence-labeled minor groove binder probes (FAM or HEX, Eurofins) are described in Supplementary file 7. Probes were designed to specifically detect the K13 M579I or C580Y propeller mutations. The efficiency and sensitivity of the TaqMan primers was assessed using standard curves comprising 10-fold serially diluted templates ranging from 10 ng to 0.001 ng. Robustness was demonstrated by high efficiency (88-95%) and R^2^ values (0.98-1.00). The quantitative accuracy in genotype calling was assessed by performing multiplex qPCR assays using mixtures of WT and mutant plasmids in fixed ratios (0:100, 20:80, 40:60, 50:50, 60:40, 80:20, 100:0). Triplicate data points clustered tightly, indicating high reproducibility in the data across the fitted curve (R^2^ = 0.89 to 0.91).

Purified gDNA from fitness co-cultures was subsequently amplified and labeled using the primers and probes described in Supplementary file 7. qPCR reactions for each sample were run in triplicate. 20 μL reactions consisted of 1×QuantiFAST reaction mix containing ROX reference dye (Qiagen, Germany), 0.66 µM of forward and reverse primers, 0.16 µM FAM-MGB and HEX-MGB TaqMan probes, and 10 ng genomic DNA. Amplification and detection of fluorescence were carried out on the QuantStudio 3 (Applied Biosystems) using the genotyping assay mode. Cycling conditions were as follows: 30s at 60°C; 5 min at 95°C; and 40 cycles of 30s at 95°C and 1 min at 60°C for primer annealing and extension. Every assay was run with positive controls (WT or mutant plasmids at different fixed ratios). No-template negative controls (water) in triplicates were processed in parallel. Rn, the fluorescence of the FAM or HEX probe, was normalized to the fluorescence signal of the ROX reporter dye. Background-normalized fluorescence (Rn minus baseline, or ΔRn) was calculated as a function of cycle number.

To determine the WT or mutant allele frequency in each sample, we first confirmed the presence of the allele by only retaining values where the threshold cycle (C_t_) of the sample was less than the no-template control by at least three cycles. Next, we subtracted the ΔRn of the samples from the background ΔRn of the no-template negative control. We subsequently normalized the fluorescence to 100% using the positive control plasmids to obtain the percentage of the WT and mutant alleles for each sample. The final percentage of the mutant allele was defined as the average of these two values: the normalized percentage of the mutant allele, and 100% minus the normalized percentage of the wild-type allele.

### eGFP-based fitness assays

Fitness assays with Dd2 parasite lines were performed as previously described (Ross *et al*., 2018). Briefly, *K13*-edited parasite lines were co-cultured in 1:1 ratios with an eGFP-positive (eGFP^+^) Dd2 reporter line. Fitness assays were initiated with tightly synchronized trophozoites in 96-well plates, with 200 μL culture volumes. Percentages of eGFP^+^ parasites were monitored by flow cytometry every two days over a period of 20 days (10 generations). Flow cytometry was performed as written above, except that only 100 nM MitoTracker DeepRed staining was used to detect total parasitemias, since SYBR Green and eGFP fluoresce in the same channel.

### Fitness Costs

The fitness cost associated with a line expressing a given K13 mutation was calculated was calculated relative to its isogenic WT counterpart using the following equation:

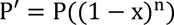

where P’ is equal to the parasitemia at the assay endpoint, P is equal to the parasitemia on day 0, n is equal to the number of generations from the assay start to finish, and x is equal to the fitness cost. This equation assumes 100% growth for the WT comparator line. For qPCR and GFP-based fitness assays, days 32 and 20 were set as the assay endpoints, resulting in the number of parasite generations (n) being set to 16 and 10, respectively.

### Ethics statement

Health care facilities were in charge of collecting anonymized *P. falciparum* positive cases. Identification of individuals cannot be established. The studies were approved by ethics committees listed in Supplementary file 4. We note that the sponsors had no role in the study design or in the collection or analysis of the data. There was no confidentiality agreement between the sponsors and the investigators.

## Acknowledgments

We thank Dr. Pascal Ringwald (World Health Organization) for his support and feedback. DAF gratefully acknowledges the US National Institutes of Health (R01 AI109023), the Department of Defense (W81XWH1910086) and the Bill & Melinda Gates Foundation (OPP1201387) for their financial support. BHS was funded in part by T32 AI106711 (PD: D. Fidock). SM is a recipient of a Human Frontiers of Science Program Long-Term Fellowship. CHC was supported in part by the NIH (R01 AI121558; PI: Jonathan Juliano). FN is supported by the Wellcome Trust of Great Britain (Grant ID: 106698). TJCA acknowledges funding support from the NIH (R37 AI048071). DAF and DM gratefully acknowledge the World Health Organization for their funding. We thank the following individuals for their kind help with the *K13*-genotyped samples – Chad: Ali S. Djiddi, Mahamat S. I. Diar, Kodbessé Boulotigam, Mbanga Djimadoum, Hamit M. Alio, Mahamat M. H. Taisso, Issa A. Haggar; Burkina Faso: TES 2017-2018 team and the US President’s Malaria Initiative through the Improving Malaria Care Project as the funding agency for the study in Burkina Faso, Chris-Boris G. Panté-Wockama; Burundi: Dismas Baza; Tanzania: Mwaka Kakolwa, Celine Mandara, Tanzania TES coordination team for the Ministry of Health; Sierra Leone: Anitta R. Y. Kamara, Foday Sahr, Mohamed Samai; The Gambia: Balla Kandeh, Joseph Okebe, Serign J. Ceesay, Baboucarr Babou, Emily Jagne, Alsan Jobe; Congo: Brice S. Pembet, Jean M. Youndouka; Somalia: Jamal Ghilan Hefzullah Amran, Abdillahi Mohamed Hassan, Abdikarim Hussein Hassan and Ali Abdulrahman; Rwanda: extended TES team for the Malaria and Other Parasitic Diseases Division, Rwanda Biomedical Centre.

## Supplementary Information

**Figure 1–source data 1.**
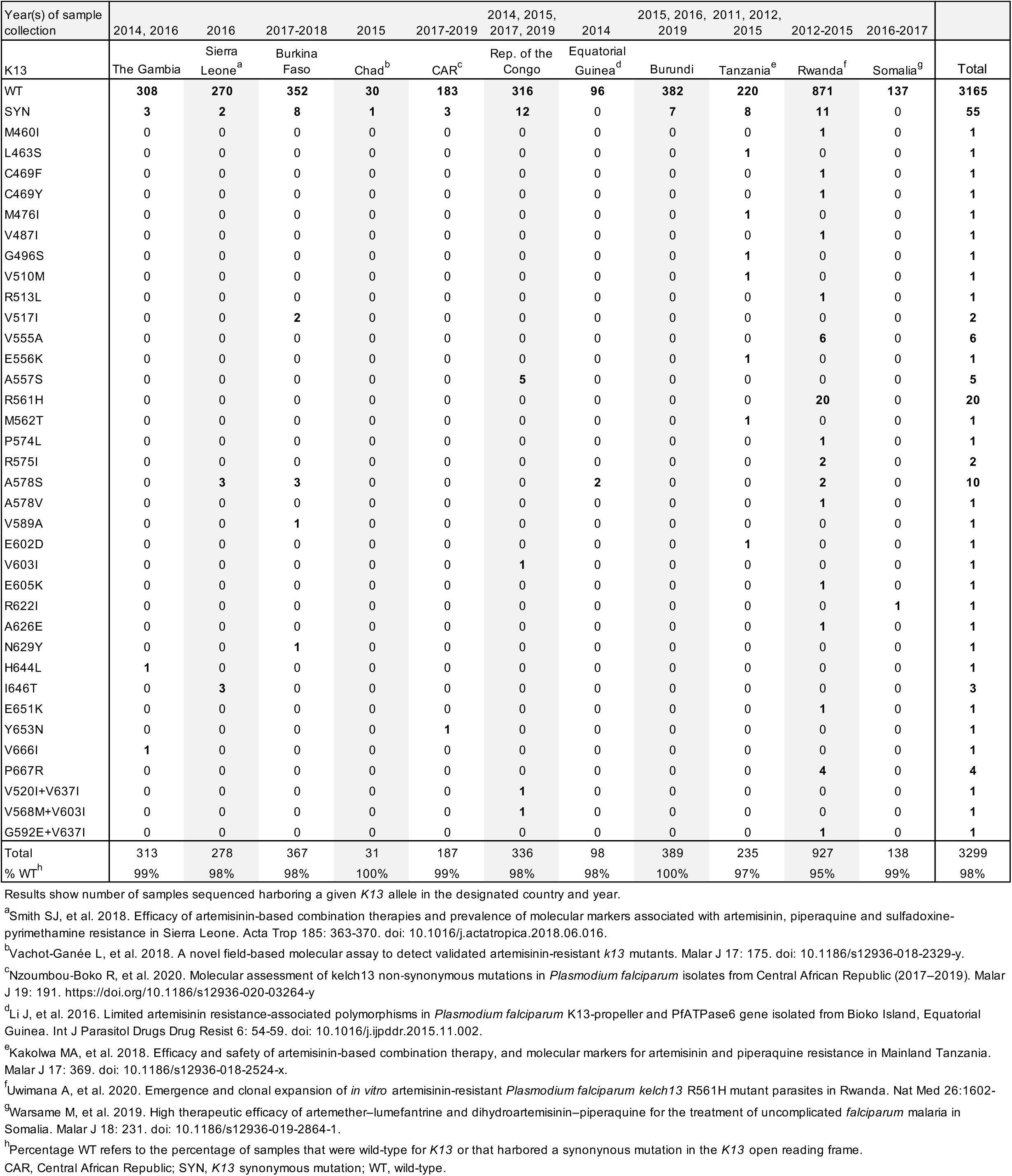
Distribution of *K13* alleles over time in African countries.

**Table 1–table supplement 1.**
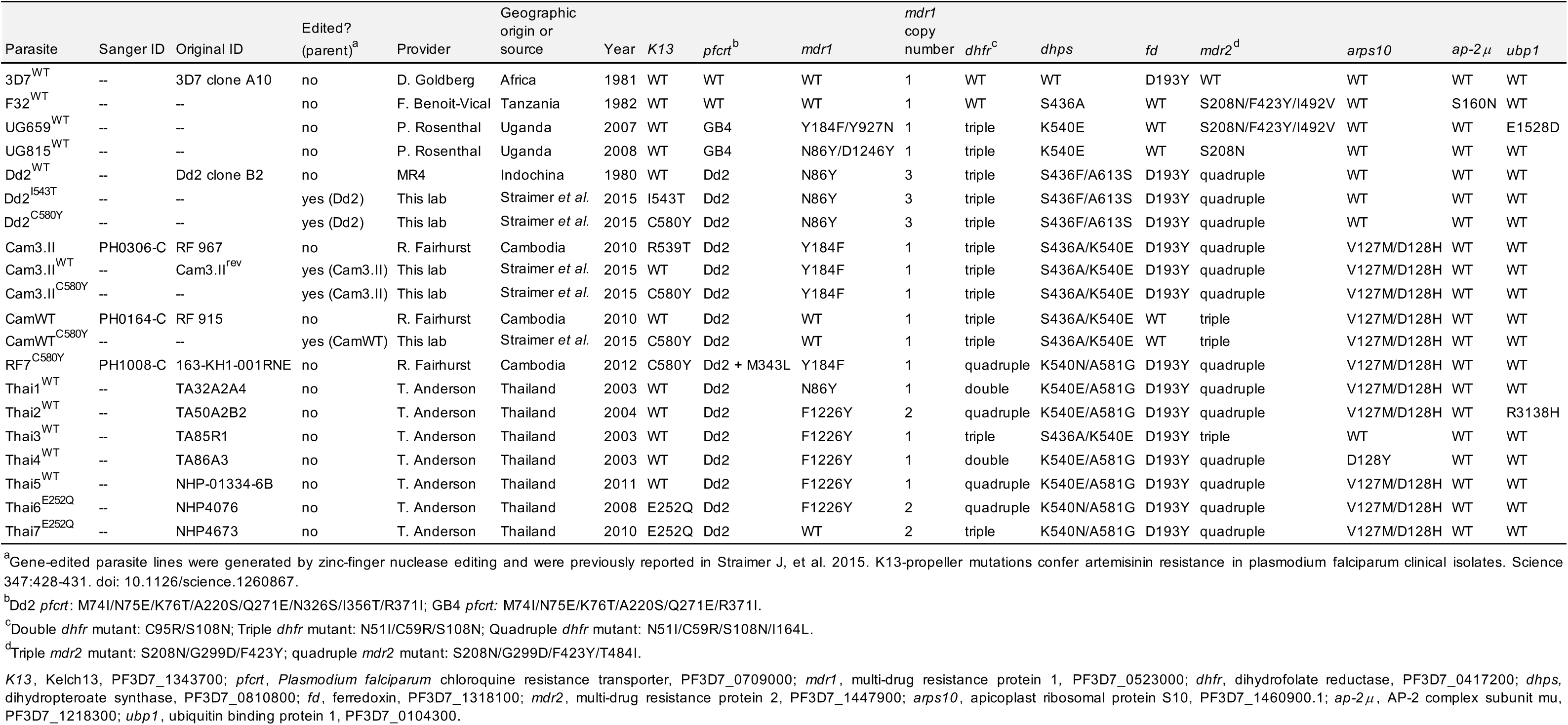
Geographic origin and drug resistance genotypes of Plasmodium falciparum clinical isolates and reference lines employed in this study.

**Figure 2–source data 1.** Ring-stage survival (RSA) assay data for edited parasites and controls (African strains).

| Parasite | Parent | K13 | Editing method | RSA survival<sup>a</sup> |  |  |  |
| --- | --- | --- | --- | --- | --- | --- | --- |
|  |  |  |  | Mean | SEM | N<sup>b</sup> | P value<sup>c</sup> |
| 3D7<sup>WT</sup> | -- | WT | -- | 1.4 | 0.3 | 4 | -- |
| 3D7<sup>ctrl</sup> | 3D7 | WT + bsm | CRISPR-Cas9 | 1.3 | 0.2 | 3 | 0.78 (ns) |
| 3D7<sup>R561H</sup> | 3D7 | R561H | CRISPR-Cas9 | 6.6 | 0.3 | 3 | <0.0001 (****) |
| 3D7<sup>M579I</sup> | 3D7 | M579I | CRISPR-Cas9 | 4.8 | 0.6 | 3 | 0.0023 (**) |
| 3D7<sup>C580Y</sup> | 3D7 | C580Y | CRISPR-Cas9 | 4.8 | 0.4 | 4 | 0.0006 (***) |
| F32<sup>WT</sup> | -- | WT | -- | 0.3 | 0.04 | 7 | -- |
| F32<sup>R561H</sup> | F32 | R561H | CRISPR-Cas9 | 0.4 | 0.16 | 3 | 0.21 (ns) |
| F32<sup>M579I</sup> | F32 | M579I | CRISPR-Cas9 | 0.5 | 0.09 | 6 | 0.043 (*) |
| F32<sup>C580Y</sup> | F32 | C580Y | CRISPR-Cas9 | 0.3 | 0.04 | 8 | 0.93 |
| UG659<sup>WT</sup> | -- | WT | -- | 1.0 | 0.1 | 6 | -- |
| UG659<sup>M579I</sup> | UG659 | M579I | CRISPR-Cas9 | 6.3 | 0.7 | 6 | <0.0001 (****) |
| UG659<sup>C580Y</sup> | UG659 | C580Y | CRISPR-Cas9 | 4.7 | 1.0 | 6 | 0.0046 (**) |
| UG815<sup>WT</sup> | -- | WT | -- | 1.3 | 0.2 | 7 | -- |
| UG815<sup>M579I</sup> clone 1 | UG815 | M579I | CRISPR-Cas9 | 11.9 | 1.0 | 7 | <0.0001 (****) |
| UG815<sup>M579I</sup> clone 2 | UG815 | M579I | CRISPR-Cas9 | 11.8 | 0.8 | 7 | <0.0001 (****) |
| UG815<sup>C580Y</sup> | UG815 | C580Y | CRISPR-Cas9 | 11.8 | 1.5 | 7 | <0.0001 (****) |
| Dd2<sup>WT</sup> | -- | WT | -- | 0.6 | 0.1 | 13 | -- |
| Dd2<sup>M579I</sup> | Dd2 | M579I | CRISPR-Cas9 | 4.0 | 0.5 | 10 | <0.0001 (****) |
| Dd2<sup>C580Y</sup> | Dd2 | C580Y | ZFN | 4.7 | 0.4 | 9 | <0.0001 (****) |

<sup>a</sup>RSA survival values represent the percentage of parasites surviving a 6 h pulse of 700nM dihydroartemisinin, calculated relative to DMSO mock-treated controls. Parasitemias were determined 66 h following drug treatment.
<sup>b</sup>N, number of independent experiments, each with technical duplicates.
<sup>c</sup>P value calculated for relative to the respective wild-type parental line. P value determined by unpaired *t* test. ns, not significant. \* *P*<0.05; \*\* *P*<0.01, \*\*\* *P*<0.001, \*\*\*\* *P*<0.0001.
bsm, binding-site mutations; SEM, standard error of the mean; WT, wild-type.

**Figure 3–source data 1.** Fitness assay data (fraction of K13 mutant parasites of total WT and mutant) for edited African parasite lines.

| Day | 3D7 vs. 3D7<sup>M579I</sup> |  |  | 3D7 vs. 3D7<sup>C580Y</sup> |  |  | F32 vs. 3D7<sup>M579I</sup> |  |  | F32 vs. F32<sup>C580Y</sup> |  |  | UG659 vs. UG659<sup>M579I</sup> |  |  | UG659 vs. UG659<sup>C580Y</sup> |  |  | UG815 vs. UG815<sup>M579I</sup> |  |  | UG815 vs. UG815<sup>C580Y</sup> |  |  |
| --- | --- | --- | --- | --- | --- | --- | --- | --- | --- | --- | --- | --- | --- | --- | --- | --- | --- | --- | --- | --- | --- | --- | --- | --- |
|  | Mean<sup>a</sup> | SEM | N<sup>b</sup> | Mean | SEM | N | Mean | SEM | N | Mean | SEM | N | Mean | SEM | N | Mean | SEM | N | Mean | SEM | N | Mean | SEM | N |
| 0<sup>c</sup> | 0.50 | 0.00 | 5 | 0.50 | 0.00 | 5 | 0.50 | 0.00 | 5 | 0.50 | 0.00 | 5 | 0.50 | 0.00 | 2 | 0.50 | 0.00 | 2 | 0.50 | 0.00 | 2 | 0.50 | 0.00 | 2 |
| 4 | 0.40 | 0.04 | 5 | 0.36 | 0.04 | 5 | 0.49 | 0.01 | 5 | 0.60 | 0.08 | 5 | 0.44 | 0.01 | 2 | 0.51 | 0.07 | 2 | 0.27 | 0.04 | 2 | 0.55 | 0.21 | 2 |
| 8 | 0.54 | 0.12 | 5 | 0.52 | 0.13 | 5 | 0.47 | 0.03 | 5 | 0.49 | 0.10 | 5 | 0.43 | 0.04 | 2 | 0.50 | 0.05 | 2 | 0.19 | 0.00 | 2 | 0.40 | 0.16 | 2 |
| 12 | 0.34 | 0.04 | 5 | 0.33 | 0.05 | 5 | 0.44 | 0.01 | 5 | 0.47 | 0.04 | 5 | 0.41 | 0.01 | 2 | 0.49 | 0.07 | 2 | 0.18 | 0.01 | 2 | 0.29 | 0.17 | 2 |
| 16 | 0.28 | 0.05 | 5 | 0.31 | 0.03 | 5 | 0.42 | 0.01 | 5 | 0.41 | 0.05 | 5 | 0.36 | 0.03 | 2 | 0.47 | 0.05 | 2 | 0.15 | 0.07 | 2 | 0.18 | 0.14 | 2 |
| 20 | 0.29 | 0.05 | 5 | 0.36 | 0.06 | 5 | 0.34 | 0.04 | 5 | 0.35 | 0.06 | 5 | 0.34 | 0.00 | 2 | 0.43 | 0.02 | 2 | 0.11 | 0.06 | 2 | 0.24 | n/a | 1 |
| 24 | 0.19 | 0.06 | 5 | 0.23 | 0.04 | 5 | 0.30 | 0.03 | 5 | 0.36 | 0.06 | 5 | 0.31 | 0.02 | 2 | 0.41 | 0.04 | 2 | 0.10 | 0.06 | 2 | 0.21 | n/a | 1 |
| 28 | 0.17 | 0.09 | 5 | 0.28 | 0.06 | 5 | 0.29 | 0.03 | 5 | 0.36 | 0.06 | 5 | 0.26 | 0.00 | 2 | 0.40 | 0.04 | 2 | 0.09 | 0.01 | 2 | 0.18 | n/a | 1 |
| 32 | 0.17 | 0.07 | 5 | 0.17 | 0.02 | 5 | 0.26 | 0.04 | 5 | 0.34 | 0.07 | 5 | 0.19 | 0.01 | 2 | 0.38 | 0.08 | 2 | 0.07 | 0.04 | 2 | 0.18 | n/a | 1 |
| 36 | 0.17 | 0.07 | 4 | 0.11 | 0.04 | 4 | 0.28 | 0.04 | 4 | 0.36 | 0.08 | 4 | 0.2192 | n/a | 1 | 0.42 | 0.00 | 1 | 0.05 | 0.02 | 2 | 0.09 | 0.11 | 2 |
| 40 | 0.15 | 0.09 | 4 | 0.10 | 0.01 | 4 | 0.19 | 0.07 | 4 | 0.25 | 0.10 | 3 | 0.1867 | n/a | 1 | 0.43 | 0.00 | 1 | 0.05 | 0.05 | 2 | 0.09 | 0.08 | 2 |

<sup>a</sup>Fraction of K13 mutant parasites determined by allelic discrimination via TaqMan qPCR.
<sup>b</sup>N, number of independent experiments, each performed in duplicate.
<sup>c</sup>Percent mutant parasites normalized to 50% on Day 0 for all assays.
n/a, not applicable. SEM, standard error of the mean.

**Figure 4–source data 1.** Distribution of *K13* alleles over time in Cambodia (2001-2017).

| Years of sample collection | 2001-02 |  |  |  | 2004-05 |  |  |  | 2006-07 |  |  |  | 2008-09 |  |  |  | 2010-11 |  |  |  | 2012-13 |  |  |  | 2014-15 |  |  |  | 2016-17 |  |  |  |  |
| --- | --- | --- | --- | --- | --- | --- | --- | --- | --- | --- | --- | --- | --- | --- | --- | --- | --- | --- | --- | --- | --- | --- | --- | --- | --- | --- | --- | --- | --- | --- | --- | --- | --- |
| *K13* | W | N | E | S | W | N | E | S | W | N | E | S | W | N | E | S | W | N | E | S | W | N | E | S | W | N | E | S | W | N | E | S | Total |
| WT | 58 | 25 | 66 | n/a | 30 | 27 | 30 | n/a | 9 | 48 | 22 | n/a | 19 | n/a | n/a | n/a | 46 | 110 | 104 | 4 | 12 | 59 | 225 | 13 | 18 | 31 | 74 | 2 | 0 | 68 | 93 | 1 | 1194 |
| G449A | 0 | 0 | 0 | n/a | 0 | 0 | 0 | n/a | 0 | 0 | 0 | n/a | 2 | n/a | n/a | n/a | 0 | 0 | 0 | 0 | 0 | 0 | 0 | 0 | 0 | 0 | 0 | 0 | 0 | 0 | 0 | 0 | 2 |
| N458Y | 0 | 0 | 0 | n/a | 0 | 0 | 0 | n/a | 0 | 0 | 0 | n/a | 0 | n/a | n/a | n/a | 2 | 0 | 0 | 1 | 0 | 0 | 0 | 0 | 0 | 0 | 0 | 0 | 0 | 0 | 0 | 0 | 3 |
| C469F | 0 | 0 | 0 | n/a | 0 | 0 | 0 | n/a | 0 | 0 | 0 | n/a | 0 | n/a | n/a | n/a | 0 | 0 | 0 | 0 | 0 | 0 | 0 | 0 | 0 | 0 | 1 | 0 | 0 | 0 | 0 | 0 | 1 |
| T474I | 0 | 0 | 1 | n/a | 0 | 0 | 0 | n/a | 0 | 1 | 0 | n/a | 0 | n/a | n/a | n/a | 0 | 0 | 0 | 0 | 0 | 0 | 0 | 0 | 0 | 0 | 0 | 0 | 0 | 0 | 0 | 0 | 2 |
| A481V | 0 | 0 | 0 | n/a | 1 | 0 | 0 | n/a | 1 | 0 | 0 | n/a | 2 | n/a | n/a | n/a | 0 | 0 | 0 | 0 | 0 | 0 | 0 | 0 | 0 | 0 | 0 | 0 | 0 | 0 | 0 | 0 | 4 |
| N489D | 0 | 0 | 0 | n/a | 0 | 0 | 0 | n/a | 0 | 0 | 0 | n/a | 0 | n/a | n/a | n/a | 0 | 0 | 0 | 0 | 0 | 0 | 0 | 0 | 0 | 2 | 0 | 0 | 0 | 0 | 0 | 0 | 2 |
| Y493H | 6 | 0 | 0 | n/a | 0 | 0 | 0 | n/a | 9 | 0 | 0 | n/a | 17 | n/a | n/a | n/a | 18 | 21 | 1 | 0 | 0 | 5 | 0 | 0 | 5 | 1 | 0 | 0 | 0 | 0 | 0 | 0 | 83 |
| I543T | 0 | 0 | 0 | n/a | 0 | 0 | 0 | n/a | 0 | 0 | 0 | n/a | 0 | n/a | n/a | n/a | 1 | 0 | 3 | 0 | 0 | 0 | 0 | 0 | 0 | 0 | 0 | 0 | 0 | 0 | 0 | 0 | 4 |
| G533S | 0 | 0 | 1 | n/a | 0 | 0 | 0 | n/a | 0 | 0 | 0 | n/a | 0 | n/a | n/a | n/a | 0 | 0 | 0 | 0 | 0 | 0 | 0 | 0 | 0 | 0 | 0 | 0 | 0 | 0 | 0 | 0 | 1 |
| N537D | 0 | 0 | 0 | n/a | 0 | 0 | 0 | n/a | 0 | 0 | 0 | n/a | 0 | n/a | n/a | n/a | 0 | 0 | 0 | 0 | 0 | 0 | 0 | 0 | 0 | 1 | 0 | 0 | 0 | 0 | 0 | 0 | 1 |
| R539T | 8 | 1 | 0 | n/a | 5 | 0 | 0 | n/a | 7 | 0 | 0 | n/a | 18 | n/a | n/a | n/a | 15 | 21 | 0 | 2 | 1 | 6 | 1 | 0 | 1 | 0 | 0 | 0 | 0 | 0 | 1 | 0 | 87 |
| P553L | 0 | 0 | 0 | n/a | 0 | 0 | 0 | n/a | 0 | 0 | 0 | n/a | 0 | n/a | n/a | n/a | 0 | 0 | 0 | 1 | 0 | 0 | 1 | 0 | 0 | 0 | 2 | 0 | 0 | 0 | 1 | 0 | 5 |
| R561H | 8 | 0 | 0 | n/a | 1 | 0 | 0 | n/a | 0 | 0 | 0 | n/a | 0 | n/a | n/a | n/a | 0 | 0 | 0 | 0 | 0 | 0 | 0 | 0 | 0 | 0 | 0 | 0 | 0 | 0 | 0 | 0 | 9 |
| V568G | 0 | 0 | 0 | n/a | 0 | 0 | 0 | n/a | 0 | 0 | 0 | n/a | 0 | n/a | n/a | n/a | 0 | 0 | 2 | 0 | 0 | 0 | 0 | 0 | 0 | 0 | 0 | 0 | 0 | 0 | 0 | 0 | 2 |
| P574L | 0 | 0 | 1 | n/a | 3 | 0 | 0 | n/a | 0 | 0 | 0 | n/a | 0 | n/a | n/a | n/a | 0 | 0 | 0 | 0 | 0 | 0 | 0 | 0 | 0 | 0 | 0 | 0 | 0 | 0 | 0 | 0 | 4 |
| C580Y | 23 | 0 | 0 | n/a | 41 | 0 | 0 | n/a | 39 | 0 | 0 | n/a | 47 | n/a | n/a | n/a | 191 | 90 | 50 | 11 | 180 | 42 | 16 | 37 | 189 | 156 | 71 | 7 | 33 | 207 | 405 | 80 | 1915 |
| D584V | 0 | 0 | 0 | n/a | 0 | 0 | 0 | n/a | 0 | 0 | 0 | n/a | 4 | n/a | n/a | n/a | 0 | 0 | 0 | 0 | 0 | 0 | 1 | 0 | 0 | 0 | 0 | 0 | 0 | 0 | 0 | 0 | 5 |
| K610R | 0 | 0 | 0 | n/a | 0 | 0 | 0 | n/a | 0 | 0 | 0 | n/a | 0 | n/a | n/a | n/a | 0 | 0 | 0 | 0 | 0 | 0 | 0 | 0 | 0 | 1 | 0 | 0 | 0 | 1 | 0 | 0 | 2 |
| A626E | 0 | 0 | 0 | n/a | 0 | 0 | 0 | n/a | 0 | 0 | 0 | n/a | 0 | n/a | n/a | n/a | 0 | 0 | 0 | 0 | 0 | 0 | 0 | 0 | 0 | 0 | 1 | 0 | 0 | 0 | 0 | 0 | 1 |
| Total | 103 | 26 | 69 | n/a | 81 | 27 | 30 | n/a | 65 | 49 | 22 | n/a | 109 | n/a | n/a | n/a | 273 | 242 | 160 | 19 | 193 | 112 | 244 | 50 | 213 | 192 | 149 | 9 | 33 | 276 | 500 | 81 | 3327 |
| % WT | 56% | 96% | 96% | n/a | 37% | 100% | 100% | n/a | 14% | 98% | 100% | n/a | 17% | n/a | n/a | n/a | 17% | 45% | 65% | 21% | 6% | 53% | 92% | 26% | 8% | 16% | 50% | 22% | 0% | 25% | 19% | 1% | 36% |
| % C580Y | 22% | 0% | 0% | n/a | 51% | 0% | 0% | n/a | 60% | 0% | 0% | n/a | 43% | n/a | n/a | n/a | 70% | 37% | 31% | 58% | 93% | 38% | 7% | 74% | 89% | 81% | 48% | 78% | 100% | 75% | 81% | 99% | 58% |

Results show number of samples sequenced harboring a given *K13* allele in each of four regions of Cambodia during a given span of time. W, Western; N, Northern; E, Eastern; S, Southern.
n/a, not available; WT, wild-type.

**Fgure 4–figure supplement 1.**
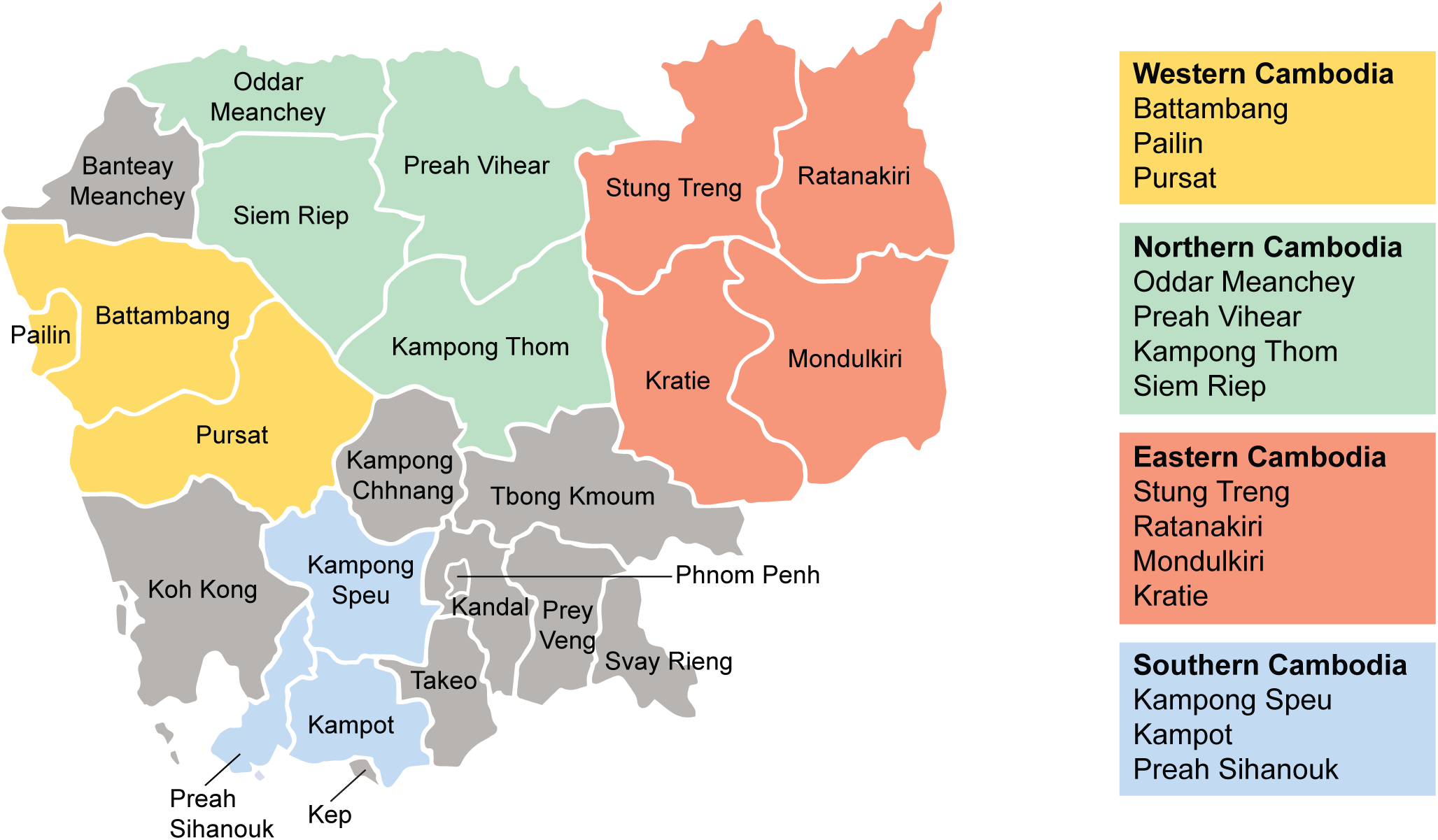
Regions of sample collection in Cambodia for K13 sequencing. Map depicting the four regions of Cambodia (Western, Northern, Eastern, and Southern) in which samples were collected between 2001 and 2017 for K13 genotyping. Genotyping data are presented in Figure 4 and are tabulated in Figure 4–source data 1.

**Figure 5–source data 1.** Ring-stage survival (RSA) assay data for edited parasites and controls (Southeast Asian strains).

| Parasite | Parent | K13 | Editing method | RSA survival<sup>a</sup> |  |  |  |
| --- | --- | --- | --- | --- | --- | --- | --- |
|  |  |  |  | Mean | SEM | N<sup>b</sup> | P value<sup>c</sup> |
| Dd2<sup>WT</sup> | — | WT | — | 0.6 | 0.1 | 13 | — |
| Dd2<sup>ctrl</sup> | Dd2 | WT + bsm<sup>d</sup> | CRISPR-Cas9 | 0.9 | 0.1 | 5 | 0.16 (ns) |
| Dd2<sup>E252Q</sup> | Dd2 | E252Q | ZFN | 0.5 | 0.2 | 5 | 0.47 (ns) |
| Dd2<sup>F446I</sup> | Dd2 | F446I | CRISPR-Cas9 | 2.0 | 0.4 | 6 | 0.0004 (***) |
| Dd2<sup>P553L</sup> | Dd2 | P553L | CRISPR-Cas9 | 4.6 | 0.7 | 11 | <0.0001 (****) |
| Dd2<sup>R561H</sup> | Dd2 | R561H | CRISPR-Cas9 | 4.3 | 0.4 | 7 | <0.0001 (****) |
| Dd2<sup>P574L</sup> | Dd2 | P574L | CRISPR-Cas9 | 2.1 | 0.3 | 8 | <0.0001 (****) |
| Dd2<sup>C580Y</sup> | Dd2 | C580Y | ZFN | 4.7 | 0.4 | 9 | <0.0001 (****) |
| Dd2<sup>R539T</sup> | Dd2 | R539T | ZFN | 20.0 |  | 1 | <0.0001 (****) |
| Cam3.II<sup>WT</sup> | Cam3.II | WT + bsm<sup>e</sup> | ZFN | 1.5 | 0.5 | 5 | — |
| Cam3.II<sup>ctrl</sup> | Cam3.II<sup>WT</sup> | WT + bsm<sup>d</sup> | CRISPR-Cas9 | 1.9 | 0.1 | 2 | 0.72 (ns) |
| Cam3.II<sup>E252Q</sup> | Cam3.II<sup>WT</sup> | E252Q | CRISPR-Cas9 | 1.0 | 0.1 |  | 0.58 (ns) |
| Cam3.II<sup>F446I</sup> | Cam3.II<sup>WT</sup> | F446I | CRISPR-Cas9 | 1.9 | 0.8 | 4 | 0.73 (ns) |
| Cam3.II<sup>P553L</sup> | Cam3.II<sup>WT</sup> | P553L | CRISPR-Cas9 | 2.8 | 0.4 | 3 | 0.14 (ns) |
| Cam3.II<sup>P574L</sup> | Cam3.II<sup>WT</sup> | P574L | CRISPR-Cas9 | 2.7 | 1.2 | 5 | 0.38 (ns) |
| Cam3.II<sup>R561H</sup> | Cam3.II<sup>WT</sup> | R561H | CRISPR-Cas9 | 13.2 | 1.3 | 6 | <0.0001 (****) |
| Cam3.II<sup>C580Y</sup> | Cam3.II | C580Y | ZFN | 10.0 |  | 1 | <0.0001 (****) |
| Cam3.II<sup>R539T</sup> | — | R539T | — | 20.4 | 1.3 | 3 | <0.0001 (****) |

<sup>a</sup>RSA survival values represent the percentage of parasites surviving a 6 h pulse of 700nM dihydroartemisinin, calculated relative to DMSO mock-treated controls. Parasitemias were determined 66 h following drug treatment.
<sup>b</sup>N, number of independent experiments, each with technical duplicates.
<sup>c</sup>P value calculated for relative to the respective wild-type parental line. P value determined by unpaired *t* test. ns, not significant. \*\*\* *P*<0.001, \*\*\*\* *P*<0.0001.
<sup>d</sup>The Dd2<sup>ctrl</sup> and Cam3.II<sup>ctrl</sup> lines harbor silent binding-site mutations at the CRISPR-Cas9 cleavage site.
<sup>e</sup>The Cam3.II<sup>WT</sup> line harbors silent binding-site mutations at the zinc-finger nuclease cleavage site.
bsm, binding-site mutations; SEM, standard error of the mean; WT, wild-type; ZFN, zinc-finger nuclease.

**Figure 5–source data 2.** Fitness assay data (percent eGFP+ parasites) for edited Dd2 parasites and parental control.

| Day | Dd2<sup>WT</sup> |  |  | Dd2<sup>bsm</sup> |  |  | Dd2<sup>E252Q</sup> |  |  | Dd2<sup>F446I</sup> |  |  | Dd2<sup>P553L</sup> |  |  | Dd2<sup>R561H</sup> |  |  | Dd2<sup>P574L</sup> |  |  | Dd2<sup>C580Y</sup> |  |  |
| --- | --- | --- | --- | --- | --- | --- | --- | --- | --- | --- | --- | --- | --- | --- | --- | --- | --- | --- | --- | --- | --- | --- | --- | --- |
|  | Mean<sup>a</sup> | SEM | N<sup>b</sup> | Mean | SEM | N | Mean | SEM | N | Mean | SEM | N | Mean | SEM | N | Mean | SEM | N | Mean | SEM | N | Mean | SEM | N |
| 0 | 48.9 | 1.4 | 3 | 49.9 | 1.3 | 3 | 48.2 | 4.2 | 3 | 50.2 | 1.2 | 3 | 50.8 | 3.6 | 3 | 47.1 | 3.0 | 3 | 45.2 | 0.7 | 3 | 50.3 | 1.4 | 3 |
| 2 | 40.0 | 3.7 | 3 | 41.9 | 2.8 | 3 | 43.8 | 1.9 | 3 | 42.6 | 7.6 | 3 | 41.6 | 4.5 | 3 | 43.0 | 4.5 | 3 | 39.5 | 5.8 | 3 | 46.2 | 4.3 | 3 |
| 4 | 34.6 | 0.7 | 2 | 38.6 | 1.3 | 2 | 44.5 | 4.5 | 2 | 37.0 | 4.4 | 2 | 39.0 | 2.4 | 2 | 39.4 | 1.5 | 2 | 34.9 | 5.3 | 2 | 42.8 | 0.3 | 2 |
| 6 | 32.1 | 0.3 | 2 | 38.2 | 1.4 | 2 | 45.0 | 4.2 | 2 | 36.3 | 3.7 | 2 | 37.6 | 0.3 | 2 | 39.6 | 2.3 | 2 | 33.6 | 3.0 | 2 | 39.2 | 3.2 | 2 |
| 8 | 25.5 | 1.3 | 2 | 33.1 | 0.3 | 2 | 40.5 | 6.8 | 2 | 32.5 | 0.3 | 2 | 27.1 | 0.7 | 2 | 35.9 | 4.4 | 2 | 29.4 | 1.8 | 2 | 30.7 | 3.6 | 2 |
| 10 | 22.4 | — | 1 | 32.0 | — | 1 | 28.8 | — | 1 | 28.5 | — | 1 | 23.5 | — | 1 | 33.7 | — | 1 | 24.8 | — | 1 | 24.5 | — | 1 |
| 12 | 15.3 | 1.1 | 2 | 24.8 | 2.1 | 2 | 25.8 | 0.3 | 2 | 22.0 | 1.5 | 2 | 20.1 | 0.2 | 2 | 27.3 | 1.1 | 2 | 19.1 | 0.5 | 2 | 18.0 | 0.8 | 2 |
| 14 | 14.3 | 1.9 | 3 | 21.6 | 0.5 | 3 | 28.3 | 4.6 | 3 | 18.8 | 0.7 | 3 | 16.3 | 1.7 | 3 | 25.1 | 0.4 | 3 | 15.7 | 1.6 | 3 | 17.1 | 2.0 | 3 |
| 16 | 14.1 | 3.2 | 3 | 24.6 | 2.1 | 3 | 29.5 | 5.5 | 3 | 19.6 | 2.5 | 3 | 17.3 | 3.5 | 3 | 26.5 | 4.1 | 3 | 17.1 | 2.4 | 3 | 15.4 | 4.5 | 3 |
| 18 | 8.9 | 1.1 | 2 | 16.8 | 1.8 | 2 | 23.0 | 5.8 | 2 | 14.0 | 1.2 | 2 | 11.5 | 2.1 | 2 | 21.4 | 3.9 | 2 | 10.6 | 1.3 | 2 | 11.7 | 2.7 | 2 |
| 20 | 7.0 | 1.0 | 3 | 13.9 | 1.8 | 3 | 17.7 | 3.5 | 3 | 11.0 | 1.9 | 3 | 9.0 | 1.6 | 3 | 15.7 | 2.9 | 3 | 8.5 | 2.5 | 3 | 6.6 | 1.4 | 3 |

<sup>a</sup>Percent eGFP+ parasites determined by flow cytometry as a fraction of total (MitoTracker+) parasite population.
<sup>b</sup>N, number of independent experiments, each performed in triplicate.
bsm; binding-site mutations; SEM, standard error of the mean; WT, wild-type.

**Figure 5–figure supplement 1.**
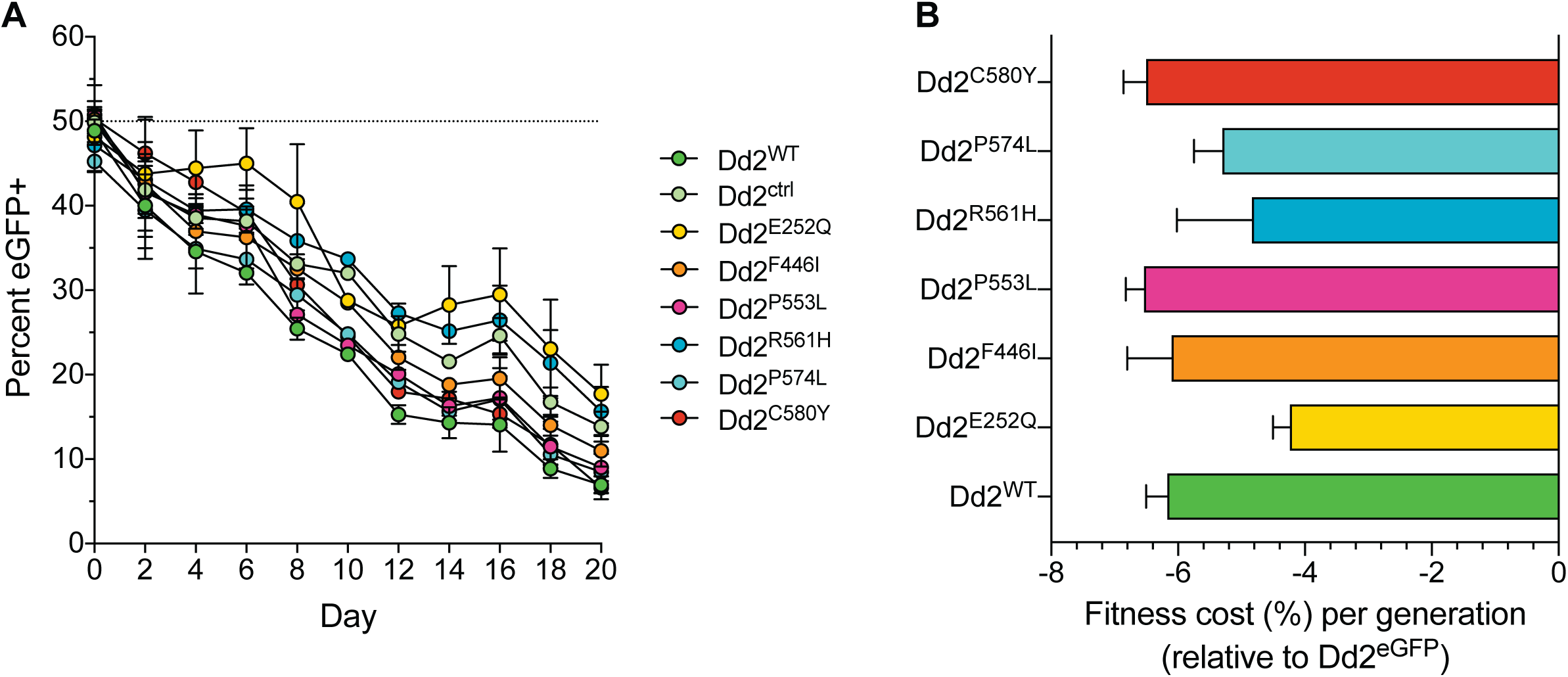
Southeast Asian K13 mutations result in minor *in vitro* growth defects in Dd2 parasites, with the exception of the C580Y and P553L mutations. (A) Percentage of eGFP+ parasites over time in parasite cultures in which an eGFP-expressing Dd2 line was co-cultured in a 1:1 mixture with either the Dd2 K13 WT parental line (Dd2^WT^) or individual Dd2 gene-edited K13 mutant lines. Parasite lines were co-cultured over a period of 20 days and percentages of eGFP+ parasites were determined by flow cytometry. Data are shown as means ± SEM (detailed in Figure 5–source data 2). Results were obtained from three independent experiments, each performed in triplicate. (B) Percent reduction in growth rate per 48 h generation, termed the fitness cost, are shown as mean ± SEM values for each mutant line relative to the Dd2^eGFP^ line.

**Figure 6–source data 1.**
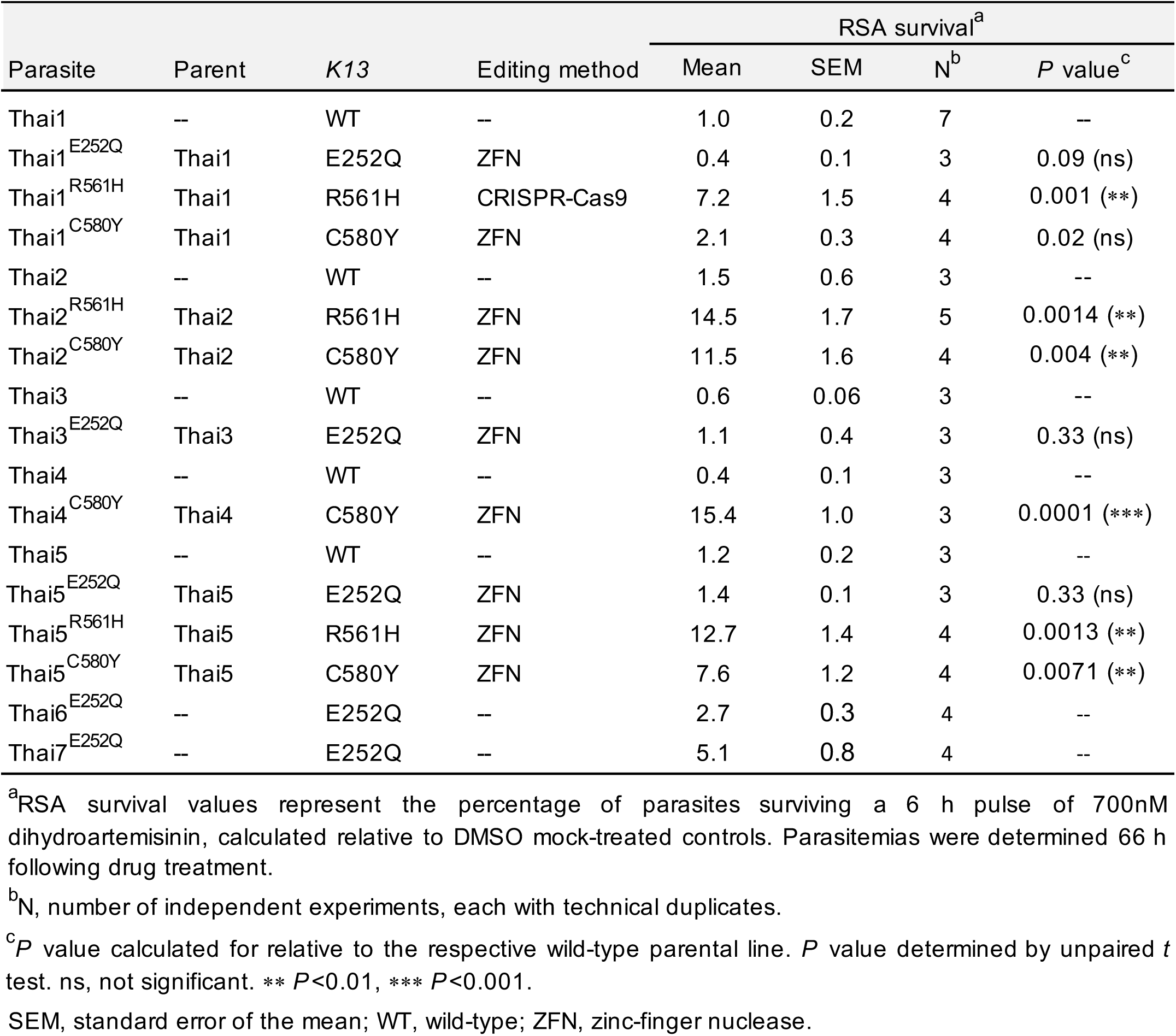
Ring-stage survival (RSA) assay data for edited parasites and controls (Thai strains).

**Figure 7–source data 1.**
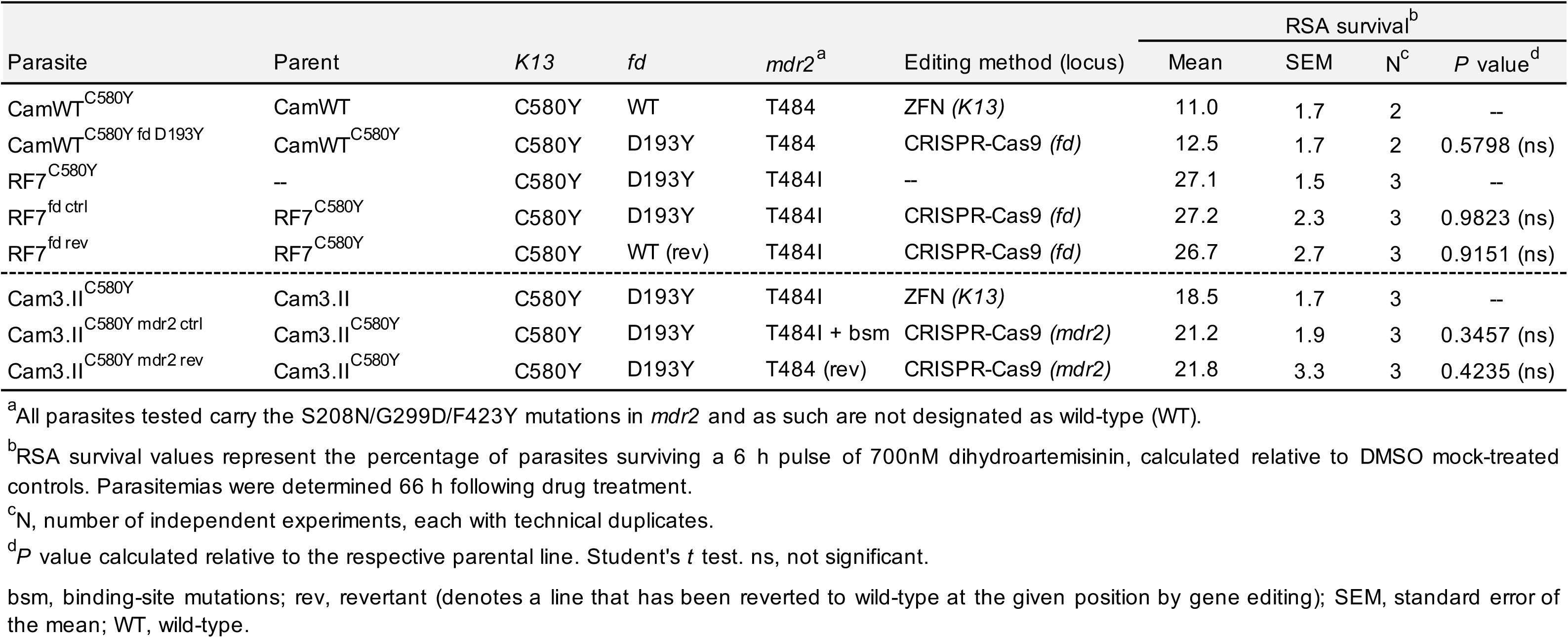
Ring-stage survival (RSA) assay data for *fd* and *mdr2* edited parasites and parental controls.

| Parasite | Parent | *K13* | *fd* | *mdr2*<sup>a</sup> | Editing method (locus) | RSA survival<sup>b</sup> |  |  |  |
| --- | --- | --- | --- | --- | --- | --- | --- | --- | --- |
|  |  |  |  |  |  | Mean | SEM | N<sup>c</sup> | *P* value<sup>d</sup> |
| CamWT<sup>C580Y</sup> | CamWT | C580Y | WT | T484 | ZFN (*K13*) | 11.0 | 1.7 | 2 | -- |
| CamWT<sup>C580Y fd D193Y</sup> | CamWT<sup>C580Y</sup> | C580Y | D193Y | T484 | CRISPR-Cas9 (*fd*) | 12.5 | 1.7 | 2 | 0.5798 (ns) |
| RF7<sup>C580Y</sup> | -- | C580Y | D193Y | T484I | -- | 27.1 | 1.5 | 3 | -- |
| RF7<sup>fd ctrl</sup> | RF7<sup>C580Y</sup> | C580Y | D193Y | T484I | CRISPR-Cas9 (*fd*) | 27.2 | 2.3 | 3 | 0.9823 (ns) |
| RF7<sup>fd rev</sup> | RF7<sup>C580Y</sup> | C580Y | WT (rev) | T484I | CRISPR-Cas9 (*fd*) | 26.7 | 2.7 | 3 | 0.9151 (ns) |
| Cam3.II<sup>C580Y</sup> | Cam3.II | C580Y | D193Y | T484I | ZFN (*K13*) | 18.5 | 1.7 | 3 | -- |
| Cam3.II<sup>C580Y mdr2 ctrl</sup> | Cam3.II<sup>C580Y</sup> | C580Y | D193Y | T484I + bsm | CRISPR-Cas9 (*mdr2*) | 21.2 | 1.9 | 3 | 0.3457 (ns) |
| Cam3.II<sup>C580Y mdr2 rev</sup> | Cam3.II<sup>C580Y</sup> | C580Y | D193Y | T484 (rev) | CRISPR-Cas9 (*mdr2*) | 21.8 | 3.3 | 3 | 0.4235 (ns) |

<sup>a</sup>All parasites tested carry the S208N/G299D/F423Y mutations in *mdr2* and as such are not designated as wild-type (WT).
<sup>b</sup>RSA survival values represent the percentage of parasites surviving a 6 h pulse of 700nM dihydroartemisinin, calculated relative to DMSO mock-treated controls. Parasitemias were determined 66 h following drug treatment.
<sup>c</sup>N, number of independent experiments, each with technical duplicates.
<sup>d</sup>*P* value calculated relative to the respective parental line. Student's *t* test. ns, not significant.
bsm, binding-site mutations; rev, revertant (denotes a line that has been reverted to wild-type at the given position by gene editing); SEM, standard error of the mean; WT, wild-type.

**Supplementary file 1.**
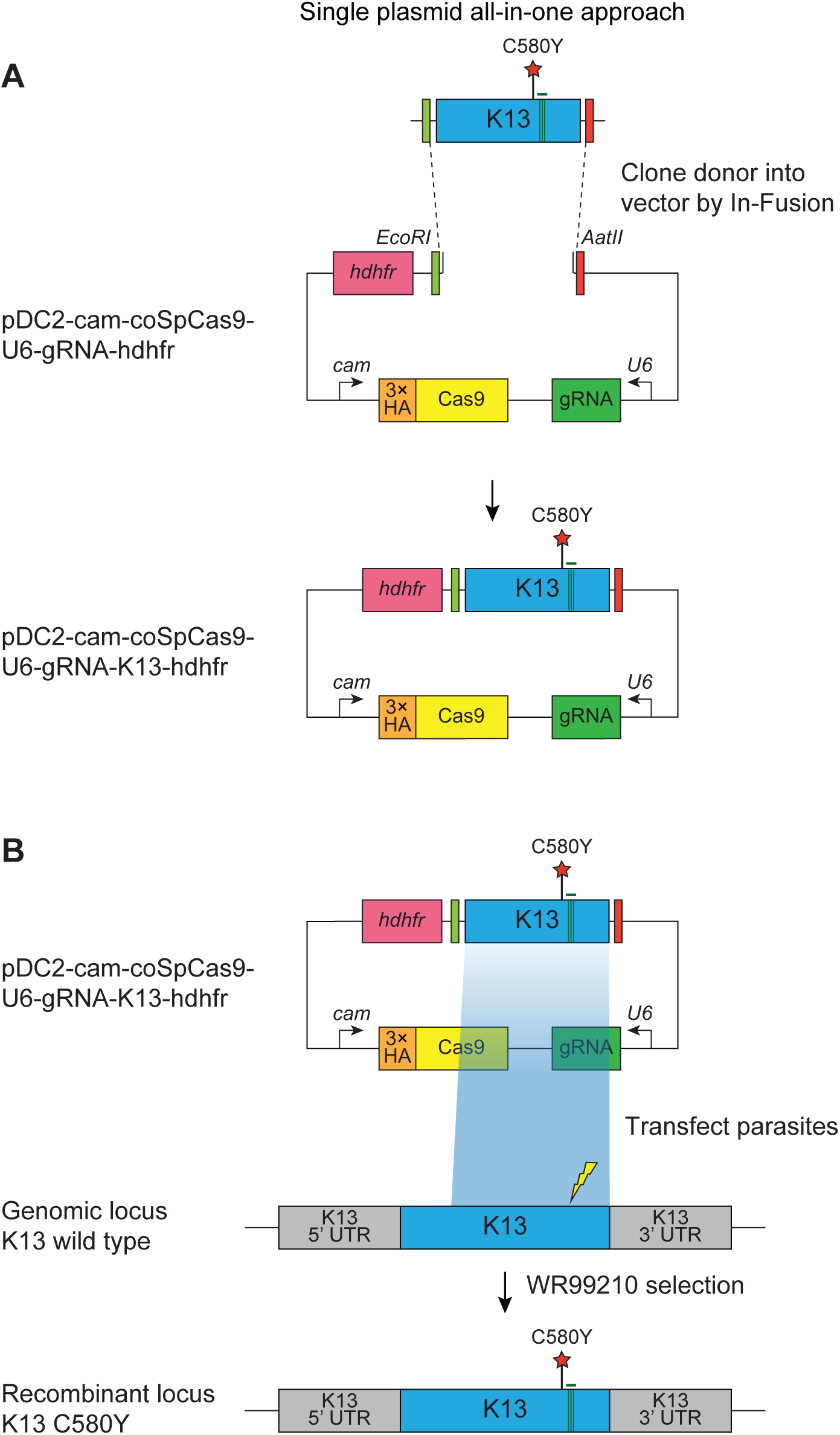
CRISPR/Cas9 strategy for editing the *K13* locus. All-in-one plasmid approach used for CRISPR/Cas9-mediated *K13* gene editing, consisting of a *K13*-specific donor template for homology-directed repair, a *K13*-specific gRNA expressed from the U6 promoter, a Cas9 cassette with expression driven by the calmodulin (cam) promoter, and a selectable marker (human *dhfr*, conferring resistance to the antimalarial WR99210 that inhibits *P. falciparum* dhfr). The Cas9 sequence was codon-optimized for improved expression in *P. falciparum*. Donors coding for specific mutations of interest (e.g. K13 C580Y, red star) were generated by site-directed mutagenesis (SDM) of the *K13* wild-type donor sequence. Green bars indicate the presence of silent binding-site mutations that were introduced by SDM to protect the edited locus from further cleavage. The lightning bolt indicates the location of the cut site in the genomic target locus. Primers used for SDM and cloning and transfection plasmids are described in Supplementary files 5 and 6, resp1e4ctively.

**Supplementary file 2.**
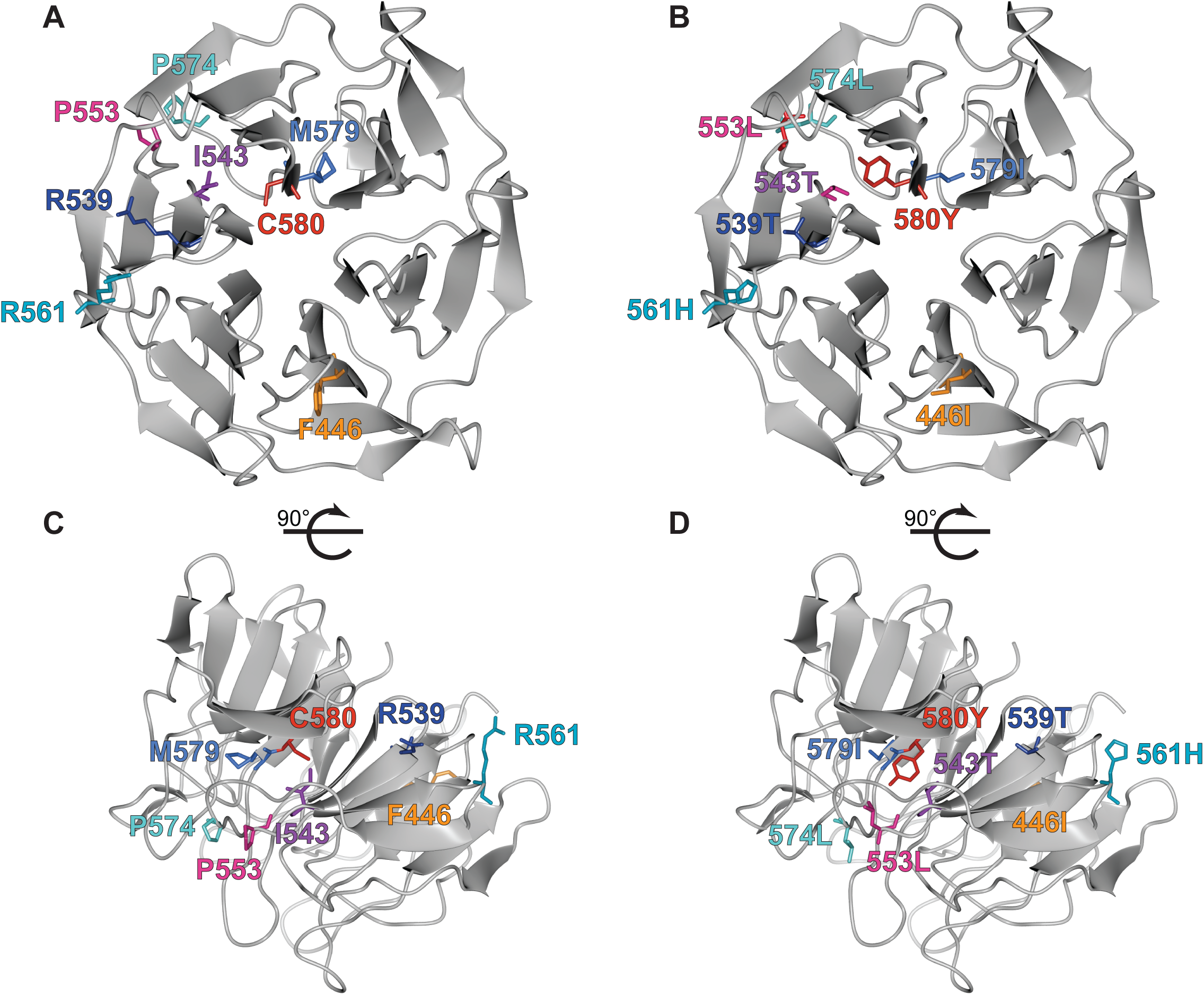
Crystal structure of K13 propeller domain showing positions of mutated residues. (A, B) Top and (C, D) side views of the crystal structure of the K13 propeller domain (PDB ID: 4YY8), highlighting residues of interest (F446I, orange; R539T, dark blue; I543T, purple; P553L, pink; R561H, dark turquoise; P574L, light turquoise; M579I medium blue; C580Y, red). Structures shown in (A) and (C) show wild-type residues while (B) and (D) show mutated residues.

**Supplementary file 3.**
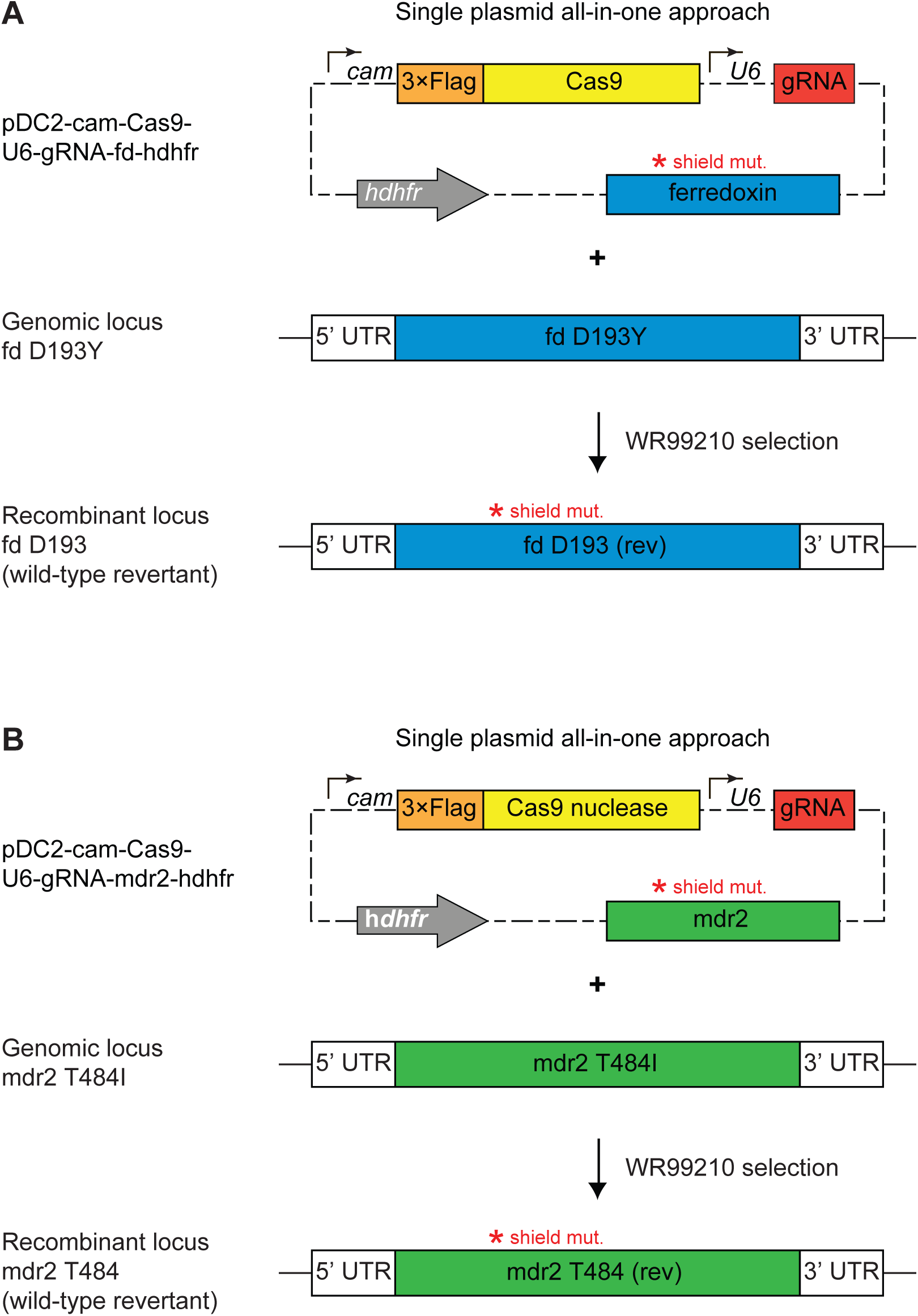
CRISPR/Cas9 strategy for editing the ferrodoxin (fd) and multidrug resis-tance protein 2 (mdr2) loci. All-in-one plasmid approaches used for CRISPR/Cas9-mediated editing of (A) the ferredoxin (fd) locus or (B) the multidrug resistance protein 2 (mdr2) locus. Plasmids consisted of a fd (A) or mdr2 (B) specific donor template for homology-directed repair, a locus-specific gRNA expressed from the U6 promoter, a Cas9 cassette with expression driven by the cam promoter, and a selectable marker (*hdhfr*, conferring resistance to WR99210). Donors coding for specific mutations of interest (i.e. fd D193Y or mdr2 T484I) were generated by site-directed mutagenesis (SDM) of the WT donor sequences. Red stars indicate the presence of silent binding-site mutations (also introduced by SDM) used to protect edited loci from further cleavage. Primers used for SDM and cloning and transfection plasmids are described in Supplementary files 5 and 6, respectively.

**Supplementary file 4.**
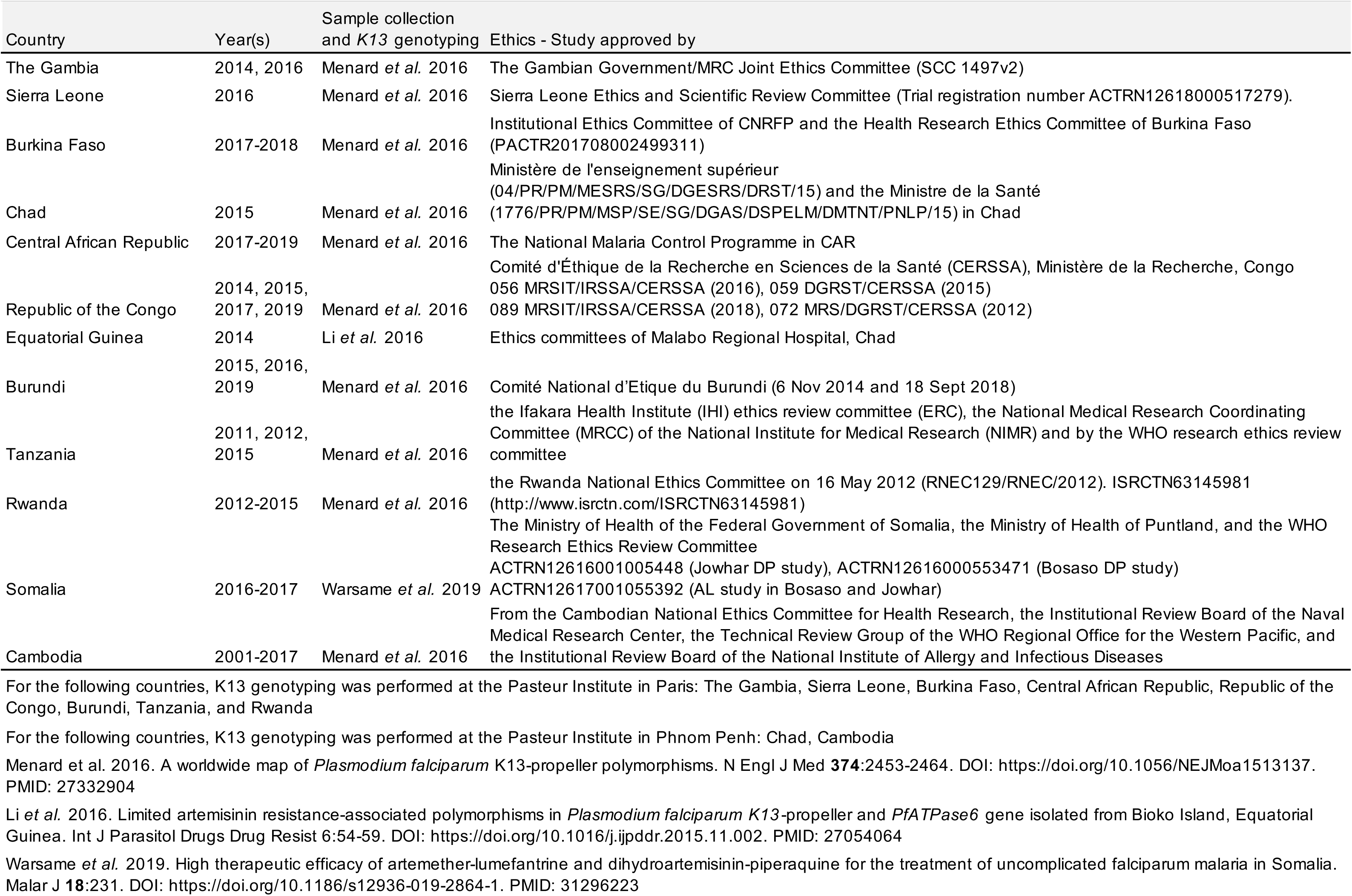
Sample collection, k13 genotyping and appoval from within-country ethics committees.

**Supplementary file 5.**
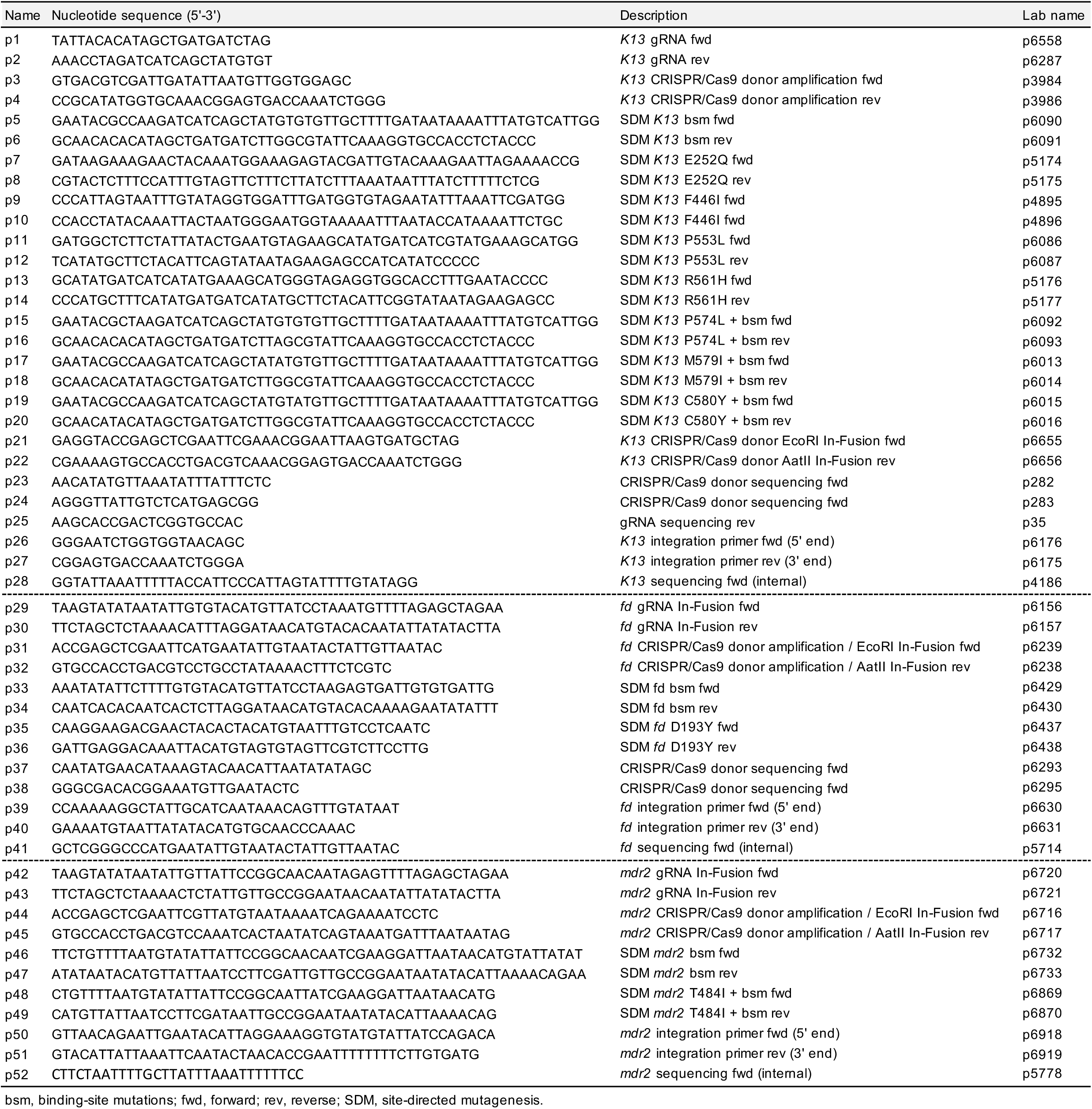
Oligonucleotides used in this study.

**Supplementary file 6.** Description of gene-editing plasmids generated in this study.

| Name | Locus | Mutations | Editing method | Lab name |
| --- | --- | --- | --- | --- |
| pDC2-cam-coSpCas9-U6-gRNA-K13_bsm-hdhfr | *K13* | *K13* bsm | CRISPR/Cas9 | gs3297 |
| pDC2-cam-coSpCas9-U6-gRNA-K13_F446I-hdhfr | *K13* | *K13* bsm + F446I | CRISPR/Cas9 | gs3298 |
| pDC2-cam-coSpCas9-U6-gRNA-K13_P553L-hdhfr | *K13* | *K13* bsm + P553L | CRISPR/Cas9 | gs3272 |
| pDC2-cam-coSpCas9-U6-gRNA-K13_R561H-hdhfr | *K13* | *K13* bsm + R561H | CRISPR/Cas9 | gs3512 |
| pDC2-cam-coSpCas9-U6-gRNA-K13_P574L-hdhfr | *K13* | *K13* bsm + P574L | CRISPR/Cas9 | gs3273 |
| pDC2-cam-coSpCas9-U6-gRNA-K13_M579I-hdhfr | *K13* | *K13* bsm + M579I<sup>a</sup> | CRISPR/Cas9 | gs3264 |
| pDC2-cam-coSpCas9-U6-gRNA-K13_C580Y-hdhfr | *K13* | *K13* bsm + C580Y | CRISPR/Cas9 | gs3262 |
| pZFN18/20-K13_E252Q-hdhfr | *K13* | *K13* bsm + E252Q | ZFN | gs3509 |
| pZFN18/20-K13_R561H-hdhfr | *K13* | *K13* bsm + R561H | ZFN | gs3510 |
| pDC2-cam-Cas9-U6-gRNA-fd_rev_D193-hdhfr | *fd* | *fd* bsm | CRISPR/Cas9 | gs3123 |
| pDC2-cam-Cas9-U6-gRNA-fd_D193Y-hdhfr | *fd* | *fd* bsm + D193Y | CRISPR/Cas9 | gs3114 |
| pDC2-cam-Cas9-U6-gRNA-mdr2_T484-hdhfr | *mdr2* | *mdr2* bsm<sup>b</sup> | CRISPR/Cas9 | gs3277 |
| pDC2-cam-Cas9-U6-gRNA-PfMdr2_T484I-hdhfr | *mdr2* | *mdr2* bsm + T484I<sup>b</sup> | CRISPR/Cas9 | gs3286 |

<sup>a</sup>*K13* donor sequence harbors both CRISPR/Cas9 and ZFN bsm in addition to the M579I mutation.
<sup>b</sup>*mdr2* donor sequence also harbors the S208N/G299D/F423Y mutations found in all parental lines.
bsm, silent binding-site mutations; ZFN, zinc-finger nuclease.

**Supplementary file 7.**
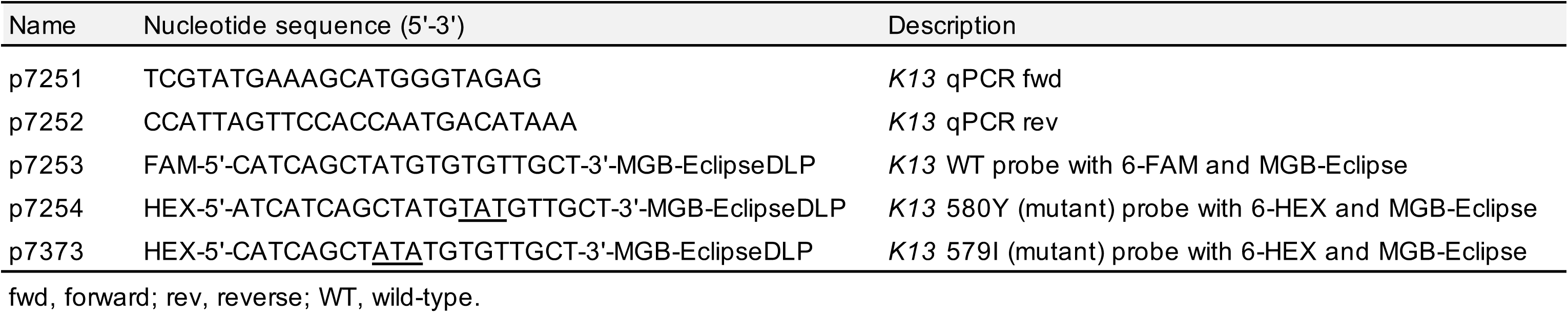
Real-Time PCR (qPCR) primers and probes.

## References

Adams T, Ennuson NAA, Quashie NB, Futagbi G, Matrevi S, Hagan OCK, Abuaku B, Koram KA, Duah NO. 2018. Prevalence of *Plasmodium falciparum* delayed clearance associated polymorphisms in adaptor protein complex 2 mu subunit (*pfap2mu*) and ubiquitin specific protease 1 (*pfubp1*) genes in Ghanaian isolates. Parasit Vectors 11:175. DOI: https://doi.org/10.1186/s13071-018-2762-3. PMID: 29530100

Amato R, Pearson RD, Almagro-Garcia J, Amaratunga C, Lim P, Suon S, Sreng S, Drury E, Stalker J, Miotto O, Fairhurst RM, Kwiatkowski DP. 2018. Origins of the current outbreak of multidrug-resistant malaria in southeast Asia: a retrospective genetic study. Lancet Infect Dis 18:337–345. DOI: https://doi.org/10.1016/S1473-3099(18)30068-9. PMID: 29398391

Anderson TJ, Nair S, McDew-White M, Cheeseman IH, Nkhoma S, Bilgic F, McGready R, Ashley E, Pyae Phyo A, White NJ, Nosten F. 2017. Population parameters underlying an ongoing soft sweep in Southeast Asian malaria parasites. Mol Biol Evol 34:131–144. DOI: https://doi.org/10.1093/molbev/msw228. PMID: 28025270

Ariey F, Witkowski B, Amaratunga C, Beghain J, Langlois AC, Khim N, Kim S, Duru V, Bouchier C, Ma L, Lim P, Leang R, Duong S, Sreng S, Suon S, Chuor CM, Bout DM, Menard S, Rogers WO, Genton B, Fandeur T, Miotto O, Ringwald P, Le Bras J, Berry A, Barale JC, Fairhurst RM, Benoit-Vical F, Mercereau-Puijalon O, Menard D. 2014. A molecular marker of artemisinin-resistant *Plasmodium falciparum* malaria. Nature 505:50–55. DOI: https://doi.org/10.1038/nature12876. PMID: 24352242

Ashley EA, Dhorda M, Fairhurst RM, Amaratunga C, Lim P, Suon S, Sreng S, Anderson JM, Mao S, Sam B, Sopha C, Chuor CM, Nguon C, Sovannaroth S, Pukrittayakamee S, Jittamala P, Chotivanich K, Chutasmit K, Suchatsoonthorn C, Runcharoen R, Hien TT, Thuy-Nhien NT, Thanh NV, Phu NH, Htut Y, Han KT, Aye KH, Mokuolu OA, Olaosebikan RR, Folaranmi OO, Mayxay M, Khanthavong M, Hongvanthong B, Newton PN, Onyamboko MA, Fanello CI, Tshefu AK, Mishra N, Valecha N, Phyo AP, Nosten F, Yi P, Tripura R, Borrmann S, Bashraheil M, Peshu J, Faiz MA, Ghose A, Hossain MA, Samad R, Rahman MR, Hasan MM, Islam A, Miotto O, Amato R, MacInnis B, Stalker J, Kwiatkowski DP, Bozdech Z, Jeeyapant A, Cheah PY, Sakulthaew T, Chalk J, Intharabut B, Silamut K, Lee SJ, Vihokhern B, Kunasol C, Imwong M, Tarning J, Taylor WJ, Yeung S, Woodrow CJ, Flegg JA, Das D, Smith J, Venkatesan M, Plowe CV, Stepniewska K, Guerin PJ, Dondorp AM, Day NP, White NJ, Tracking Resistance to Artemisinin Consortium. 2014. Spread of artemisinin resistance in Plasmodium falciparum malaria. N Engl J Med 371:411-423. DOI: https://doi.org/10.1056/NEJMoa1314981. PMID: 25075834

Asua V, Conrad MD, Aydemir O, Duvalsaint M, Legac J, Duarte E, Tumwebaze P, Chin DM, Cooper RA, Yeka A, Kamya MR, Dorsey G, Nsobya SL, Bailey J, Rosenthal PJ. 2020. Changing prevalence of potential mediators of aminoquinoline, antifolate, and artemisinin resistance across Uganda. J Infect Dis Nov 4;jiaa687. DOI: https://doi.org/10.1093/infdis/jiaa687. PMID: 33146722

Bergmann C, van Loon W, Habarugira F, Tacoli C, Jager JC, Savelsberg D, Nshimiyimana F, Rwamugema E, Mbarushimana D, Ndoli J, Sendegeya A, Bayingana C, Mockenhaupt FP. 2021. Increase in Kelch 13 polymorphisms in *Plasmodium falciparum*, Southern Rwanda. Emerg Infect Dis 27:294–296. DOI: https://doi.org/10.3201/eid2701.203527. PMID: 33350925

Birnbaum J, Scharf S, Schmidt S, Jonscher E, Hoeijmakers WAM, Flemming S, Toenhake CG, Schmitt M, Sabitzki R, Bergmann B, Frohlke U, Mesen-Ramirez P, Blancke Soares A, Herrmann H, Bartfai R, Spielmann T. 2020. A Kelch13-defined endocytosis pathway mediates artemisinin resistance in malaria parasites. Science 367:51–59. DOI: https://doi.org/10.1126/science.aax4735. PMID: 31896710

Cerqueira GC, Cheeseman IH, Schaffner SF, Nair S, McDew-White M, Phyo AP, Ashley EA, Melnikov A, Rogov P, Birren BW, Nosten F, Anderson TJC, Neafsey DE. 2017. Longitudinal genomic surveillance of *Plasmodium falciparum* malaria parasites reveals complex genomic architecture of emerging artemisinin resistance. Genome Biol 18:78. DOI: https://doi.org/10.1186/s13059-017-1204-4. PMID: 28454557

Conrad MD, Rosenthal PJ. 2019. Antimalarial drug resistance in Africa: the calm before the storm? Lancet Infect Dis 19:e338–e351. DOI: https://doi.org/10.1016/S1473-3099(19)30261-0. PMID: 31375467

Das S, Manna S, Saha B, Hati AK, Roy S. 2019. Novel *pfkelch13* gene polymorphism associates with artemisinin resistance in Eastern India. Clin Infect Dis 69:1144–1152. DOI: https://doi.org/10.1093/cid/ciy1038. PMID: 30535043

Demas AR, Sharma AI, Wong W, Early AM, Redmond S, Bopp S, Neafsey DE, Volkman SK, Hartl DL, Wirth DF. 2018. Mutations in *Plasmodium falciparum* actin-binding protein coronin confer reduced artemisinin susceptibility. Proc Natl Acad Sci USA 115:12799–12804. DOI: https://doi.org/10.1073/pnas.1812317115. PMID: 30420498

Dondorp AM, Nosten F, Yi P, Das D, Phyo AP, Tarning J, Lwin KM, Ariey F, Hanpithakpong W, Lee SJ, Ringwald P, Silamut K, Imwong M, Chotivanich K, Lim P, Herdman T, An SS, Yeung S, Singhasivanon P, Day NP, Lindegardh N, Socheat D, White NJ. 2009. Artemisinin resistance in *Plasmodium falciparum* malaria. N Engl J Med 361:455–467. DOI: https://doi.org/10.1056/NEJMoa0808859. PMID: 19641202

Frosch AE, Laufer MK, Mathanga DP, Takala-Harrison S, Skarbinski J, Claassen CW, Dzinjalamala FK, Plowe CV. 2014. Return of widespread chloroquine-sensitive *Plasmodium falciparum* to Malawi. J Infect Dis 210:1110–1114. DOI: https://doi.org/10.1093/infdis/jiu216. PMID: 24723474

Ghorbal M, Gorman M, Macpherson CR, Martins RM, Scherf A, Lopez-Rubio JJ. 2014. Genome editing in the human malaria parasite *Plasmodium falciparum* using the CRISPR-Cas9 system. Nat Biotechnol 32:819–821. DOI: https://doi.org/10.1038/nbt.2925. PMID: 24880488

Group WKG-PS. 2019. Association of mutations in the *Plasmodium falciparum* Kelch13 gene (Pf3D7_1343700) with parasite clearance rates after artemisinin-based treatments—a WWARN individual patient data meta-analysis. BMC Med 17:1. DOI: https://doi.org/10.1186/s12916-018-1207-3. PMID: 30651111

Haldar K, Bhattacharjee S, Safeukui I. 2018. Drug resistance in *Plasmodium*. Nat Rev Microbiol 16:156–170. DOI: https://doi.org/10.1038/nrmicro.2017.161. PMID: 29355852

Hanboonkunupakarn B, White NJ. 2020. Advances and roadblocks in the treatment of malaria. Br J Clin Pharmacol **Jul 12. doi:** 10.1111/bcp.14474. DOI: https://doi.org/10.1111/bcp.14474. PMID: 32656850

Henrici RC, van Schalkwyk DA, Sutherland CJ. 2019. Modification of *pfap2mu* and *pfubp1* markedly reduces ring-stage susceptibility of *Plasmodium falciparum* to artemisinin *in vitro*. Antimicrob Agents Chemother 64. DOI: https://doi.org/10.1128/AAC.01542-19. PMID: 31636063

Henriques G, Hallett RL, Beshir KB, Gadalla NB, Johnson RE, Burrow R, van Schalkwyk DA, Sawa P, Omar SA, Clark TG, Bousema T, Sutherland CJ. 2014. Directional selection at the pfmdr1, pfcrt, pfubp1, and pfap2mu loci of Plasmodium falciparum in Kenyan children treated with ACT. J Infect Dis 210:2001-2008. DOI: https://doi.org/10.1093/infdis/jiu358. PMID: 24994911

Imwong M, Dhorda M, Myo Tun K, Thu AM, Phyo AP, Proux S, Suwannasin K, Kunasol C, Srisutham S, Duanguppama J, Vongpromek R, Promnarate C, Saejeng A, Khantikul N, Sugaram R, Thanapongpichat S, Sawangjaroen N, Sutawong K, Han KT, Htut Y, Linn K, Win AA, Hlaing TM, van der Pluijm RW, Mayxay M, Pongvongsa T, Phommasone K, Tripura R, Peto TJ, von Seidlein L, Nguon C, Lek D, Chan XHS, Rekol H, Leang R, Huch C, Kwiatkowski DP, Miotto O, Ashley EA, Kyaw MP, Pukrittayakamee S, Day NPJ, Dondorp AM, Smithuis FM, Nosten FH, White NJ. 2020. Molecular epidemiology of resistance to antimalarial drugs in the Greater Mekong subregion: an observational study. Lancet Infect Dis 20:1470–1480. DOI: https://doi.org/10.1016/S1473-3099(20)30228-0. PMID: 32679084

Imwong M, Suwannasin K, Kunasol C, Sutawong K, Mayxay M, Rekol H, Smithuis FM, Hlaing TM, Tun KM, van der Pluijm RW, Tripura R, Miotto O, Menard D, Dhorda M, Day NPJ, White NJ, Dondorp AM. 2017. The spread of artemisinin-resistant *Plasmodium falciparum* in the Greater Mekong subregion: a molecular epidemiology observational study. Lancet Infect Dis 17:491–497. DOI: https://doi.org/10.1016/S1473-3099(17)30048-8. PMID: 28161569

Kayiba NK, Yobi DM, Tshibangu-Kabamba E, Tuan VP, Yamaoka Y, Devleesschauwer B, Mvumbi DM, Okitolonda Wemakoy E, De Mol P, Mvumbi GL, Hayette MP, Rosas-Aguirre A, Speybroeck N. 2020. Spatial and molecular mapping of Pfkelch13 gene polymorphism in Africa in the era of emerging *Plasmodium falciparum* resistance to artemisinin: a systematic review. Lancet Infect Dis. Oct 27;S1473-3099(20)30493-X. DOI: https://doi.org/10.1016/S1473-3099(20)30493-X. PMID: 33125913

Kublin JG, Cortese JF, Njunju EM, Mukadam RA, Wirima JJ, Kazembe PN, Djimde AA, Kouriba B, Taylor TE, Plowe CV. 2003. Reemergence of chloroquine-sensitive *Plasmodium falciparum* malaria after cessation of chloroquine use in Malawi. J Infect Dis 187:1870–1875. DOI: https://doi.org/10.1086/375419. PMID: 12792863

Laufer MK, Thesing PC, Eddington ND, Masonga R, Dzinjalamala FK, Takala SL, Taylor TE, Plowe CV. 2006. Return of chloroquine antimalarial efficacy in Malawi. N Engl J Med 355:1959–1966. DOI: https://doi.org/10.1056/NEJMoa062032. PMID: 17093247

Lim MY, LaMonte G, Lee MCS, Reimer C, Tan BH, Corey V, Tjahjadi BF, Chua A, Nachon M, Wintjens R, Gedeck P, Malleret B, Renia L, Bonamy GMC, Ho PC, Yeung BKS, Chow ED, Lim L, Fidock DA, Diagana TT, Winzeler EA, Bifani P. 2016. UDP-galactose and acetyl-CoA transporters as *Plasmodium* multidrug resistance genes. Nat Microbiol 1:16166. DOI: https://doi.org/10.1038/nmicrobiol.2016.166. PMID: 27642791

Lu F, Culleton R, Zhang M, Ramaprasad A, von Seidlein L, Zhou H, Zhu G, Tang J, Liu Y, Wang W, Cao Y, Xu S, Gu Y, Li J, Zhang C, Gao Q, Menard D, Pain A, Yang H, Zhang Q, Cao J. 2017. Emergence of indigenous artemisinin-resistant *Plasmodium falciparum* in Africa. N Engl J Med 376:991–993. DOI: https://doi.org/10.1056/NEJMc1612765. PMID: 28225668

MalariaGEN. 2016. Genomic epidemiology of artemisinin resistant malaria. eLife 5:e08714. DOI: https://doi.org/10.7554/eLife.08714. PMID: 26943619

Mathieu LC, Cox H, Early AM, Mok S, Lazrek Y, Paquet JC, Ade MP, Lucchi NW, Grant Q, Udhayakumar V, Alexandre JS, Demar M, Ringwald P, Neafsey DE, Fidock DA, Musset L. 2020. Local emergence in Amazonia of *Plasmodium falciparum* k13 C580Y mutants associated with *in vitro* artemisinin resistance. eLife 9:e51015. DOI: https://doi.org/10.7554/eLife.51015. PMID: 32394893

Menard D, Khim N, Beghain J, Adegnika AA, Shafiul-Alam M, Amodu O, Rahim-Awab G, Barnadas C, Berry A, Boum Y, Bustos MD, Cao J, Chen JH, Collet L, Cui L, Thakur GD, Dieye A, Djalle D, Dorkenoo MA, Eboumbou-Moukoko CE, Espino FE, Fandeur T, Ferreira-da-Cruz MF, Fola AA, Fuehrer HP, Hassan AM, Herrera S, Hongvanthong B, Houze S, Ibrahim ML, Jahirul-Karim M, Jiang L, Kano S, Ali-Khan W, Khanthavong M, Kremsner PG, Lacerda M, Leang R, Leelawong M, Li M, Lin K, Mazarati JB, Menard S, Morlais I, Muhindo-Mavoko H, Musset L, Na-Bangchang K, Nambozi M, Niare K, Noedl H, Ouedraogo JB, Pillai DR, Pradines B, Quang-Phuc B, Ramharter M, Randrianarivelojosia M, Sattabongkot J, Sheikh-Omar A, Silue KD, Sirima SB, Sutherland C, Syafruddin D, Tahar R, Tang LH, Toure OA, Tshibangu-wa-Tshibangu P, Vigan-Womas I, Warsame M, Wini L, Zakeri S, Kim S, Eam R, Berne L, Khean C, Chy S, Ken M, Loch K, Canier L, Duru V, Legrand E, Barale JC, Stokes B, Straimer J, Witkowski B, Fidock DA, Rogier C, Ringwald P, Ariey F, Mercereau-Puijalon O, Consortium K. 2016. A worldwide map of *Plasmodium falciparum* K13-propeller polymorphisms. N Engl J Med 374:2453–2464. DOI: https://doi.org/10.1056/NEJMoa1513137. PMID: 27332904

Miotto O, Amato R, Ashley EA, MacInnis B, Almagro-Garcia J, Amaratunga C, Lim P, Mead D, Oyola SO, Dhorda M, Imwong M, Woodrow C, Manske M, Stalker J, Drury E, Campino S, Amenga-Etego L, Thanh TN, Tran HT, Ringwald P, Bethell D, Nosten F, Phyo AP, Pukrittayakamee S, Chotivanich K, Chuor CM, Nguon C, Suon S, Sreng S, Newton PN, Mayxay M, Khanthavong M, Hongvanthong B, Htut Y, Han KT, Kyaw MP, Faiz MA, Fanello CI, Onyamboko M, Mokuolu OA, Jacob CG, Takala-Harrison S, Plowe CV, Day NP, Dondorp AM, Spencer CC, McVean G, Fairhurst RM, White NJ, Kwiatkowski DP. 2015. Genetic architecture of artemisinin-resistant *Plasmodium falciparum*. Nat Genet 47:226–234. DOI: https://doi.org/10.1038/ng.3189. PMID: 25599401

Miotto O, Sekihara M, Tachibana SI, Yamauchi M, Pearson RD, Amato R, Goncalves S, Mehra S, Noviyanti R, Marfurt J, Auburn S, Price RN, Mueller I, Ikeda M, Mori T, Hirai M, Tavul L, Hetzel MW, Laman M, Barry AE, Ringwald P, Ohashi J, Hombhanje F, Kwiatkowski DP, Mita T. 2020. Emergence of artemisinin-resistant *Plasmodium falciparum* with *kelch13* C580Y mutations on the island of New Guinea. PLoS Pathog 16:e1009133. DOI: https://doi.org/10.1371/journal.ppat.1009133. PMID: 33320907

Nair S, Li X, Arya GA, McDew-White M, Ferrari M, Nosten F, Anderson TJC. 2018. Do fitness costs explain the rapid spread of *kelch13*-C580Y substitutions conferring artemisinin resistance? Antimicrob Agents Chemother 62:e00605–00618. DOI: https://doi.org/10.1128/AAC.00605-18. PMID: 29914963

Noedl H, Se Y, Schaecher K, Smith BL, Socheat D, Fukuda MM, Artemisinin Resistance in Cambodia 1 Study Consortium. 2008. Evidence of artemisinin-resistant malaria in western Cambodia. N Engl J Med 359:2619-2620. DOI: https://doi.org/10.1056/NEJMc0805011. PMID: 19064625

Noedl H, Socheat D, Satimai W. 2009. Artemisinin-resistant malaria in Asia. N Engl J Med 361:540–541. DOI: https://doi.org/10.1056/NEJMc0900231. PMID: 19641219

Ord R, Alexander N, Dunyo S, Hallett R, Jawara M, Targett G, Drakeley CJ, Sutherland CJ. 2007. Seasonal carriage of pfcrt and pfmdr1 alleles in Gambian Plasmodium falciparum imply reduced fitness of chloroquine-resistant parasites. J Infect Dis 196:1613–1619. DOI: https://doi.org/10.1086/522154. PMID: 18008244

Phyo AP, Ashley EA, Anderson TJC, Bozdech Z, Carrara VI, Sriprawat K, Nair S, White MM, Dziekan J, Ling C, Proux S, Konghahong K, Jeeyapant A, Woodrow CJ, Imwong M, McGready R, Lwin KM, Day NPJ, White NJ, Nosten F. 2016. Declining efficacy of artemisinin combination therapy against *P. falciparum* malaria on the Thai-Myanmar border (2003-2013): The role of parasite genetic factors. Clin Infect Dis 63:784–791. DOI: https://doi.org/10.1093/cid/ciw388. PMID: 27313266

Ross LS, Dhingra SK, Mok S, Yeo T, Wicht KJ, Kumpornsin K, Takala-Harrison S, Witkowski B, Fairhurst RM, Ariey F, Menard D, Fidock DA. 2018. Emerging Southeast Asian PfCRT mutations confer *Plasmodium falciparum* resistance to the first-line antimalarial piperaquine. Nat Commun 9:3314. DOI: https://doi.org/10.1038/s41467-018-05652-0. PMID: 30115924

Sherrard-Smith E, Hogan AB, Hamlet A, Watson OJ, Whittaker C, Winskill P, Ali F, Mohammad AB, Uhomoibhi P, Maikore I, Ogbulafor N, Nikau J, Kont MD, Challenger JD, Verity R, Lambert B, Cairns M, Rao B, Baguelin M, Whittles LK, Lees JA, Bhatia S, Knock ES, Okell L, Slater HC, Ghani AC, Walker PGT, Okoko OO, Churcher TS. 2020. The potential public health consequences of COVID-19 on malaria in Africa. Nat Med 26:1411–1416. DOI: https://doi.org/10.1038/s41591-020-1025-y. PMID: 32770167

Siddiqui FA, Boonhok R, Cabrera M, Mbenda HGN, Wang M, Min H, Liang X, Qin J, Zhu X, Miao J, Cao Y, Cui L. 2020. Role of *Plasmodium falciparum* Kelch 13 protein mutations in *P. falciparum* populations from northeastern Myanmar in mediating artemisinin resistance. mBio 11:e01134–01119. DOI: https://doi.org/10.1128/mBio.01134-19. PMID: 32098812

Straimer J, Gnadig NF, Stokes BH, Ehrenberger M, Crane AA, Fidock DA. 2017. *Plasmodium falciparum* K13 mutations differentially impact ozonide susceptibility and parasite fitness *in vitro*. mBio 8:e00172–00117. DOI: https://doi.org/10.1128/mBio.00172-17. PMID: 28400526

Straimer J, Gnadig NF, Witkowski B, Amaratunga C, Duru V, Ramadani AP, Dacheux M, Khim N, Zhang L, Lam S, Gregory PD, Urnov FD, Mercereau-Puijalon O, Benoit-Vical F, Fairhurst RM, Menard D, Fidock DA. 2015. K13-propeller mutations confer artemisinin resistance in *Plasmodium falciparum* clinical isolates. Science 347:428–431. DOI: https://doi.org/10.1126/science.1260867. PMID: 25502314

Sutherland CJ, Henrici RC, Artavanis-Tsakonas K. 2020. Artemisinin susceptibility in the malaria parasite *Plasmodium falciparum*: propellers, adaptor proteins and the need for cellular healing. FEMS Microbiol Rev. Oct 23;fuaa056. DOI: https://doi.org/10.1093/femsre/fuaa056. PMID: 33095255

Uwimana A, Legrand E, Stokes BH, Ndikumana JM, Warsame M, Umulisa N, Ngamije D, Munyaneza T, Mazarati JB, Munguti K, Campagne P, Criscuolo A, Ariey F, Murindahabi M, Ringwald P, Fidock DA, Mbituyumuremyi A, Menard D. 2020. Emergence and clonal expansion of *in vitro* artemisinin-resistant *Plasmodium falciparum kelch13* R561H mutant parasites in Rwanda. Nat Med 26:1602–1608. DOI: https://doi.org/10.1038/s41591-020-1005-2. PMID: 32747827

van der Pluijm RW, Imwong M, Chau NH, Hoa NT, Thuy-Nhien NT, Thanh NV, Jittamala P, Hanboonkunupakarn B, Chutasmit K, Saelow C, Runjarern R, Kaewmok W, Tripura R, Peto TJ, Yok S, Suon S, Sreng S, Mao S, Oun S, Yen S, Amaratunga C, Lek D, Huy R, Dhorda M, Chotivanich K, Ashley EA, Mukaka M, Waithira N, Cheah PY, Maude RJ, Amato R, Pearson RD, Goncalves S, Jacob CG, Hamilton WL, Fairhurst RM, Tarning J, Winterberg M, Kwiatkowski DP, Pukrittayakamee S, Hien TT, Day NP, Miotto O, White NJ, Dondorp AM. 2019. Determinants of dihydroartemisinin-piperaquine treatment failure in *Plasmodium falciparum* malaria in Cambodia, Thailand, and Vietnam: a prospective clinical, pharmacological, and genetic study. Lancet Infect Dis 19:952–961. DOI: https://doi.org/10.1016/S1473-3099(19)30391-3. PMID: 31345710

Warsame M, Hassan AM, Hassan AH, Jibril AM, Khim N, Arale AM, Gomey AH, Nur ZS, Osman SM, Mohamed MS, Abdulrahman A, Yusuf FE, Amran JGH, Witkowski B, Ringwald P. 2019. High therapeutic efficacy of artemether-lumefantrine and dihydroartemisinin-piperaquine for the treatment of uncomplicated falciparum malaria in Somalia. Malar J 18:231. DOI: https://doi.org/10.1186/s12936-019-2864-1. PMID: 31296223

White J, Dhingra SK, Deng X, El Mazouni F, Lee MCS, Afanador GA, Lawong A, Tomchick DR, Ng CL, Bath J, Rathod PK, Fidock DA, Phillips MA. 2019. Identification and mechanistic understanding of dihydroorotate dehydrogenase point mutations in *Plasmodium falciparum* that confer *in vitro* resistance to the clinical candidate DSM265. ACS Infect Dis 5:90–101. DOI: https://doi.org/10.1021/acsinfecdis.8b00211. PMID: 30375858

WHO. 2019. World Health Organization. WHO status report on artemisinin resistance and ACT efficacy. https://www.who.int/docs/default-source/documents/publications/gmp/who-cds-gmp-2019-17-eng.pdf?ua=1.

WHO. 2020. World Health Organization. World malaria report 2020. https://www.who.int/teams/global-malaria-programme/reports/world-malaria-report-2020.

Witkowski B, Amaratunga C, Khim N, Sreng S, Chim P, Kim S, Lim P, Mao S, Sopha C, Sam B, Anderson JM, Duong S, Chuor CM, Taylor WR, Suon S, Mercereau-Puijalon O, Fairhurst RM, Menard D. 2013. Novel phenotypic assays for the detection of artemisinin-resistant Plasmodium falciparum malaria in Cambodia: in-vitro and ex-vivo drug-response studies. Lancet Infect Dis 13:1043–1049. DOI: https://doi.org/10.1016/S1473-3099(13)70252-4. PMID: 24035558

Witkowski B, Lelievre J, Barragan MJ, Laurent V, Su XZ, Berry A, Benoit-Vical F. 2010. Increased tolerance to artemisinin in *Plasmodium falciparum* is mediated by a quiescence mechanism. Antimicrob Agents Chemother 54:1872–1877. DOI: https://doi.org/10.1128/AAC.01636-09. PMID: 20160056

Xi R, Hadjipanayis AG, Luquette LJ, Kim TM, Lee E, Zhang J, Johnson MD, Muzny DM, Wheeler DA, Gibbs RA, Kucherlapati R, Park PJ. 2011. Copy number variation detection in whole-genome sequencing data using the Bayesian information criterion. Proc Natl Acad Sci USA 108:E1128–1136. DOI: https://doi.org/10.1073/pnas.1110574108. PMID: 22065754

Yang T, Yeoh LM, Tutor MV, Dixon MW, McMillan PJ, Xie SC, Bridgford JL, Gillett DL, Duffy MF, Ralph SA, McConville MJ, Tilley L, Cobbold SA. 2019. Decreased K13 abundance reduces hemoglobin catabolism and proteotoxic stress, underpinning artemisinin resistance. Cell Rep 29:2917–2928 e2915. DOI: https://doi.org/10.1016/j.celrep.2019.10.095. PMID: 31775055

